# Targeting a proteolytic neo-epitope of CUB-domain containing protein 1 in RAS-driven cancer

**DOI:** 10.1101/2021.06.14.448427

**Authors:** Shion A. Lim, Jie Zhou, Alexander J. Martinko, Yung-Hua Wang, Ekaterina V. Filippova, Veronica Steri, Donghui Wang, Soumya G. Remesh, Jia Liu, Byron Hann, Anthony A. Kossiakoff, Michael J. Evans, Kevin K. Leung, James A. Wells

**Affiliations:** Department of Pharmaceutical Chemistry, University of California San Francisco; San Francisco, USA; Department of Radiology and Biomedical Imaging, University of California San Francisco; San Francisco, USA; Helen Diller Family Comprehensive Cancer Center, University of California San Francisco; San Francisco, USA; Department of Biochemistry and Molecular Biology, The University of Chicago; Chicago, USA; Institute for Biophysical Dynamics, The University of Chicago; Chicago, USA; Preclinical Therapeutics Core, University of California San Francisco; San Francisco, USA; Chan Zuckerberg Biohub; San Francisco, USA; Department of Cellular and Molecular Pharmacology, University of California San Francisco; San Francisco, USA

## Abstract

A central challenge for any therapeutic is targeting diseased over normal cells. Proteolysis is frequently upregulated in disease and can generate proteoforms with unique neo-epitopes. We hypothesize that targeting proteolytic neo-epitopes can enable more effective and safer treatments, reflecting a conditional layer of disease-specific regulation. Here, we characterized the precise proteolytic isoforms of CUB domain containing protein 1 (CDCP1), a protein overexpressed and specifically cleaved in RAS-driven cancers. We validated that the N-terminal and C-terminal fragments of CDCP1 remain associated after proteolysis in vitro and on the surface of pancreatic cancer cells. Using a differential phage display strategy, we generated exquisitely selective recombinant antibodies that target cells harboring cleaved CDCP1 and not the full-length form using antibody-drug conjugates or a bi-specific T-cell engagers. We show tumor-specific localization and anti-tumor activity in a syngeneic pancreatic tumor model having superior safety profiles compared to a pan-CDCP1-targeting antibody. Our studies show proteolytic neo-epitopes can provide an orthogonal “AND” gate for disease-specific targeting.

**One-Sentence Summary:** Antibody-based targeting of neo-epitopes generated by disease-associated proteolysis improves the therapeutic index

## INTRODUCTION

A key to safe and effective cancer treatments is to target diseased over healthy cells. Traditionally this has been enabled by identifying targets with large expression differences between diseased vs. normal tissue, but this severely limits the druggable target space. More recently, there has been interest to target disease-associated alternate splice forms or oncogene-specific peptide-MHC complexes (*1, 2*). These too are rare events and the low abundance of peptide-MHC complexes may prevent effective targeting. Proteolysis plays critical roles in both normal and aberrant biological processes (*3, 4*), and is well-known to be upregulated and dysregulated in a variety of diseases including cancer (*5–7*). As such, there have been efforts to better understand disease-associated proteases and their substrates, and therapeutic strategies to inhibit disease-associated proteases or utilize these for conditional activation of masked therapeutics have been described (*8–12*). Tumor-associated proteolysis could also expose novel epitopes on the surface of cancer cells. We hypothesize that therapeutic agents that recognize these proteolytic neo-epitopes could address a major challenge of on-target off-tumor toxicity. However, generation of antibodies can specifically recognize proteolytically-generated neo-epitopes (*13*), particularly those on the cell surface, has not been demonstrated.

Ras activation is one of the most well-known oncogenic transformations and is implicated in many solid tumors including in nearly 90% of pancreatic cancer (*14, 15*). CUB domain containing protein 1 (CDCP1), also known as Trask, gp140, SIMA135, and CD318, is a Type I single-pass membrane protein that is highly overexpressed in a variety of solid tumors (*16–20*). CDCP1 is upregulated and functionally critical in KRAS-transformed cells and high expression of CDCP1 is associated with loss of adhesion and is linked to more aggressive, metastatic tumors and poor prognosis (*21–25*). Full-length CDCP1 is a 135-kDa protein composed of a large extracellular domain, a transmembrane domain, and a short intracellular domain, and is expressed on several types of tissue including epithelial tissue along the gastrointenstinal tract (*17, 26*). The heavily glycosylated extracellular domain of CDCP1 is predicted to contain three CUB-like domains with a β-sandwich fold, while the intracellular domain contains five tyrosine phosphorylation sites. Apart from this, the structure of CDCP1 remains unknown.

CDCP1 is proteolytically processed between the first and second CUB domains, presumably by serine proteases (*27*). Proteolysis, along with overexpression, has been associated with the tumor-promoting functions of CDCP1 (*24, 28*). CDCP1 activation leads to phosphorylation of intracellular tyrosine residues and initiation of downstream signaling pathways associated with loss of adhesion, increased migration, and anoikis (*17, 29*). There is a higher proportion of cleaved CDCP1 on the surface of more aggressive, metastatic cancer cells compared to less malignant cancer cell lines (*30, 31*). Given that protease levels and proteolytic activity are elevated in the tumor microenvironment (*5*), it is not surprising that cleaved CDCP1 is more prevalent in aggressive cancers (*31–33*), and normal tissues almost exclusively express the full-length form (*34–36*). Thus, we hypothesized that therapeutic agents that can specifically recognize and target the cleaved form of CDCP1 may expand the therapeutic window.

Here, we report the biochemical and biophysical characterization of cleaved CDCP1 and the generation of recombinant antibodies that specifically target this proteoform. We identified the exact cleavage sites of CDCP1 and found, surprisingly, that cleaved CDCP1 remains an intact complex. Using a differential phage display strategy, we selected and optimized antibodies that can specifically recognize cleaved CDCP1 with no detectable binding to uncleaved CDCP1. These antibodies selectively target cleaved CDCP1-expressing pancreatic cancer cells as antibody-drug conjugates (ADC) or bi-specific T-cell engagers. These antibodies localize to tumors harboring cleaved CDCP1 *in vivo*, reduce tumor growth, and significantly improve the safety profile compared to a pan-CDCP1-targeting antibody. Targeting proteolytic neo-epitopes offers a novel strategy to expand the therapeutic index of disease-associated antigens.

## RESULTS

### The N-terminal fragment of CDCP1 is retained upon proteolysis

To characterize CDCP1 proteolysis and target the cleaved form of CDCP1, we first generated recombinant proteins and cell lines expressing CDCP1 with an engineered cut site. CDCP1 has been reported to be cleaved at the dibasic arginine-lysine residues between the first and second CUB domains (R368/K369) (*17, 33*). We replaced this site with a PreScission Protease (Px) site to inducibly control the proteolysis of CDCP1. The ectodomain of CDCP1 with this engineered cut site was recombinantly expressed as an Fc-fusion with a C-terminal biotinylated Avi-tag (CDCP1(Px)-Fc) (**Fig 1A**). We additionally generated a variant where R368/K369 were mutated to alanine (CDCP1(R368A/K369A)-Fc) to prevent cleavage by basic residue-specific proteases, and another where only the N-terminal fragment (NTF, aa 30-369) is fused to an Fc domain (NTF-Fc) (**Fig. S1**). Efforts to recombinantly express the C-terminal ectodomain fragment (CTF, aa 370-665) alone were unsuccessful, suggesting the NTF plays a role in expression and folding. Addition of Px cleaves CDCP1(Px)-Fc into two fragments (NTF and CTF) of expected molecular weight (**Fig. 1B**). Surprisingly, we found that Px-treated CDCP1(Px)-Fc had the same size exclusion chromatography (SEC) elution profile as uncleaved CDCP1-Fc, with no evidence of an unbound NTF (**Fig. 1C**). Additionally, we observe robust binding by bio-layer interferometry (BLI) of IgG 4A06, an antibody that recognizes the NTF of CDCP1 (**Fig. 1D, Fig. S1**). To test if this was unique to Px cleavage, we generated and tested a Thrombin protease-cleavable CDCP1-Fc (CDCP1(Tx)-Fc) (**Fig. S2**) and observed the same phenomena, where the NTF does not dissociate from the CTF upon cleavage by Thrombin. To determine whether cleaved CDCP1 remains a complex on the cell membrane, we engineered HEK293T cell lines expressing the full CDCP1 protein sequence with an N-terminal FLAG-tag and the R368/K369 proteolysis site replaced with the Px recognition sequence (**Fig. 1E**). Using a commercial antibody (D1W9N) that recognizes the intracellular C-terminal region of CDCP1, we observed that addition of Px to cells cleaves CDCP1(Px) at the expected molecular weight (**Fig. 1F**). Despite this, staining by anti-FLAG and IgG 4A06 were unaffected with Px treatment, indicating that the NTF remains membrane-associated after proteolysis. Taken together, our data suggests that specific proteolysis of CDCP1 does not result in the dissociation of the NTF from the CTF.

**Figure 1:**
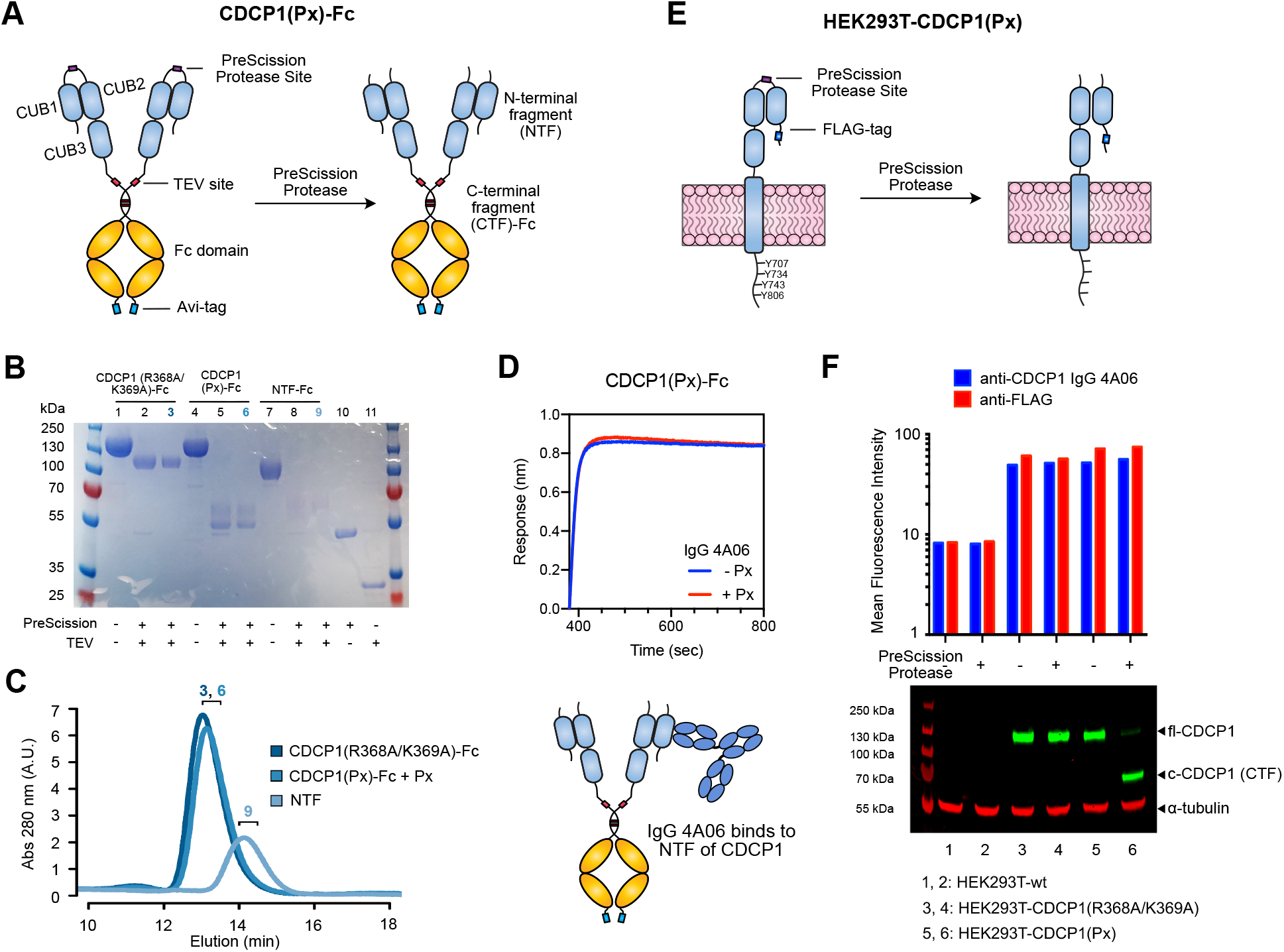
N-terminal fragment of CDCP1 is retained upon proteolysis between the CUB1 and CUB2 domain. (**A**) Design of a PreScission Protease (Px)-cleavable CDCP1 ectodomain fused to a TEV-releasable Fc domain with C-terminal Avi-tag (CDCP1(Px)-Fc). The R368/K369 cleavage site was replaced with a Px recognition sequence (GS)_5_-LEVLFQGP-(GS)_5_. (**B**) SDS-PAGE of CDCP1 constructs. Px treatment cleaves CDCP1(Px)-Fc into NTF and CTF-Fc fragments. NTF is heavily glycosylated (predicted 14 N-linked glycosylation sites) and runs as a smeared higher-molecular weight band at ∼60 kDa. (**C**) SEC traces of CDCP1(R368/K369A)-Fc and CDCP1(Px)-Fc treated with Px, and NTF (TEV released) show that the NTF and CTF of CDCP1(Px) remain intact after proteolysis. Numbers denote fractions corresponding to the SDS-PAGE gel lanes in **B**. (**D**) BLI of IgG 4A06, which recognizes the NTF, shows robust binding to both Px-treated and untreated CDCP1(Px)-Fc. (**E**) Design of Px-cleavable CDCP1 with N-terminal FLAG-tag expressed on the surface of HEK293T cells. (**F**) Flow cytometry and western blot of HEK293T-wt, HEK293T-CDCP1(R368A/K369A), HEK293T-CDCP1(Px). Flow cytometry signal of anti-FLAG and IgG 4A06 remains unchanged with Px treatment. Western blot with anti-CDCP1 D1W9N, which recognizes the C-terminal intracellular region of CDCP1, confirms Px-mediated CDCP1 proteolysis at the intended molecular weight.

### IP-MS reveals three major cleavage sites for CDCP1

Knowing the precise proteolysis site(s) of CDCP1 on cancer cells is critical to designing the appropriate antigen for antibody generation. We characterized a panel of pancreatic ductal adenocarcinoma (PDAC) cell lines (HPAC, PL5, PL45) expressing differential amounts of full-length and cleaved CDCP1 and used immunoprecipitation mass spectrometry (IP-MS) to map proteolytic peptides near the reported cut site of CDCP1 (**Fig. 2A**). HPAC primarily express uncleaved CDCP1, whereas PL5 and PL45 express cleaved CDCP1, and HPNE is a non-malignant pancreatic cell line with minimal CDCP1 expression (**Fig. 2B**). IP was performed with either D1W9N or IgG 4A06 which recognizes either the CTF or NTF, respectively. The samples were treated with Glu-C to preserve basic cut site(s) of CDCP1 and analyzed by MS. In PL5 and PL45 cells, we identified peptides mapping to both the NTF and CTF of CDCP1, regardless of which Ab was used for the IP, providing further evidence that endogenous cleaved CDCP1 is a complex on PDAC cells (**Fig. 2C, Fig. S3**). We identified three unique cut sites after basic residues between the CUB1 and CUB2 domain boundaries; these include proteolysis after K365 (Cut1), R368 (Cut2), K369 (Cut3). Cut2 and Cut3 are previously reported proteolysis sites (*37*), while Cut1 is a novel site. We observe some “frayed” peptides, suggestive of exopeptidase activity. Peptides corresponding to the uncleaved sequence were exclusively observed for HPAC, and no peptides mapping to CDCP1 were observed for HPNE reflecting the expected absence of CDCP1 expression. These findings confirm that endogenous cleaved CDCP1 remains as a complex on PDAC cells and that CDCP1 is proteolytically processed between CUB1 and CUB2 to produce a heterogenous set of cleaved forms.

**Figure 2:**
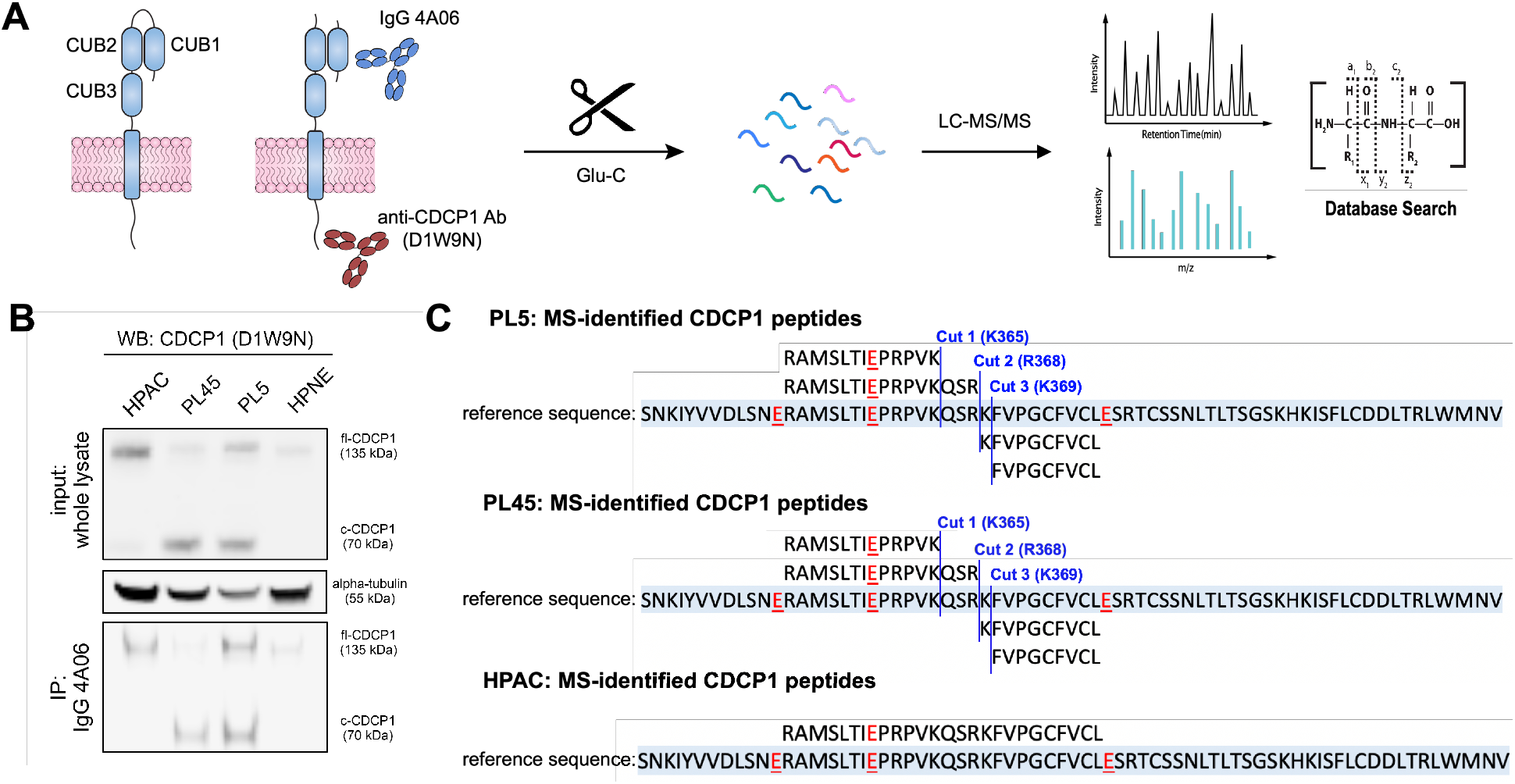
Identification of endogenous cut sites of CDCP1 on the surface of PDAC cells. (**A**) Schematic of IP-MS strategy to identify the endogenous proteolysis sites of CDCP1 on PDAC cells. CDCP1 was IP-ed with IgG 4A06 or D1W9N Ab and digested with Glu-C, which cleaves after aspartic acid, and was analyzed by LC-MS/MS to identify peptides corresponding to proteolytic products of CDCP1. (**B**) (*top*) Western blot of PDAC cell lines expressing differential amounts of uncleaved and cleaved CDCP1. D1W9N Ab was used to detect C-terminal fragment of CDCP1. PL5 and PL45 express mostly cleaved CDCP1, while HPAC expresses mostly uncleaved CDCP1. HPNE, a non-malignant pancreatic cell line, expresses low levels of CDCP1. (*bottom*) IP-blot shows that IP with IgG 4A06 can pull-down the CTF of CDCP1. (**C**) Depiction of peptides and proteolysis sites identified in PL5, PL45, and HPAC cell lines. Peptides identified by LC-MS for each cell line is aligned to the reference sequence highlighted in light blue. Aspartic acid residues recognized by Glu-C are highlighted in red underlined text. Three proteolysis sites of CDCP1: Cut 1 (K365), Cut 2 (R368), Cut 3 (K369), are observed in PL5 and PL45 cells but not on HPAC cells, and are highlighted in blue text and blue vertical lines.

### Cleaved and uncleaved CDCP1 adopt similar conformations

We hypothesized that because cleaved CDCP1 forms a tight complex, we could generate recombinant cleaved CDCP1 with the native cut sites by co-transfection of the two fragments. We designed one set of plasmids encoding the NTF of CDCP1, ending after the P1 residue of the 3 different cut sites, and a second set of plasmids encoding the CTF ectodomain of CDCP1 starting at the P1’ residue (c-CDCP1 Cut 1, Cut 2, Cut 3) (**Fig. 3A**). Uncleaved CDCP1 ectodomain was generated with a single plasmid encoding the entire ectodomain (fl-CDCP1). Indeed, after co-transfection of pairs of NTF and CTF plasmids, we were able to purify an intact cleaved CDCP1 complex for all of the cut variants (**Fig. 3B**, **Fig. S4A-D**). Both fl- and c-CDCP1 bind robustly to IgG 4A06 (**Fig. 3C**) and have similar SEC profiles (**Fig. S4E**), demonstrating that cleaved CDCP1 forms a complex even when generated *in trans*. Remarkably, these constructs all have similar melting temperatures (**Fig. 3D**), indicating that the NTF/CTF complex is stable and do not dissociate until unfolding of the entire ectodomain. We examined the secondary structure of fl- and c-CDCP1 ectodomains by circular dichroism (CD) spectroscopy (**Fig. 3E**), SEC-small-angle X-ray scattering (SEC-SAXS) (**Fig. 3F, Fig. S5**) and SEC-Multi-angle light scattering (SEC-MALS) (**Fig. 3G**). The CD spectra of fl- and c-CDCP1 show a classic β-sheet signal, consistent with the CUB domain fold. There is a noticeable change in the spectral shape and minima between fl- and c-CDCP1, which suggests that proteolysis may cause subtle changes in the secondary structure of CDCP1. Comparison of the pair distance distribution functions (P(r) functions) obtained by SEC-SAXS shows that both fl- and c-CDCP1 exhibit similar overall domain arrangement with no large-scale conformational changes as a result of proteolysis (**Fig. 3F**) (*38*). The small differences in P(r) function most likely result from subtle differences in flexibility and conformational sampling between the isoforms. fl-CDCP1 appears more flexible than c-CDCP1 as indicated by the differences in radii of gyration and small peak-shift in the dimensionless Kratky plots (**Fig. 3F**, **Fig. S5B**). Furthermore, SEC-MALS shows that fl- and c-CDCP1 have the same elution profile and molecular weight (∼97-99 kDa), consistent with the approximate size of a monomeric ectodomain (77 kDa plus glycosylation) (**Fig. 3G, Table S1**). Overall, these data show that other than small differences in the β-sheet signature, the conformation of fl- and c-CDCP1 are remarkably similar.

**Figure 3:**
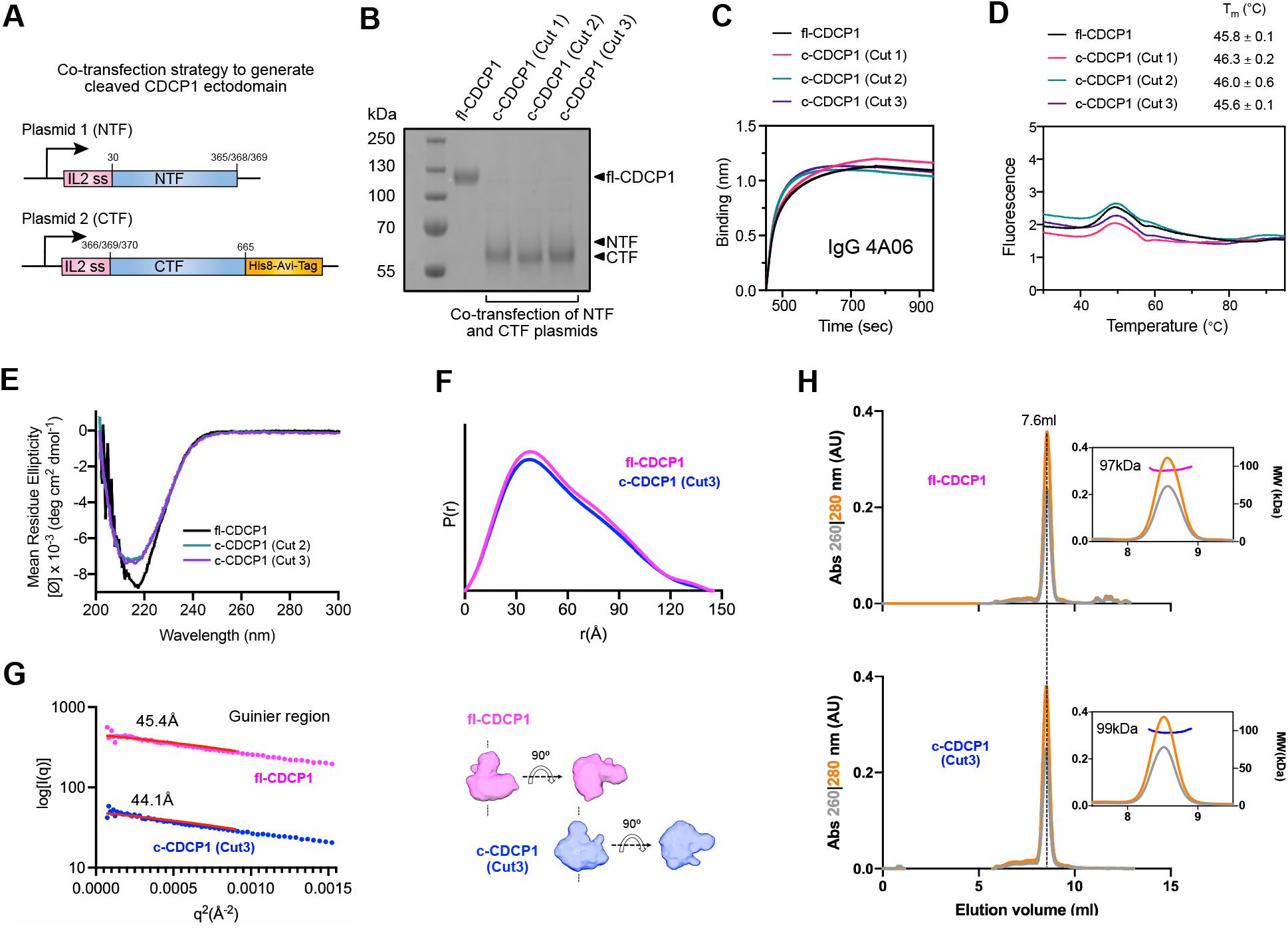
Cleaved and uncleaved CDCP1 have similar conformations. (**A**) Schematic of the co-transfection strategy to generate c-CDCP1 ectodomain. The NTF and CTF are encoded on separate plasmids with an IL2 secretion sequence. (**B**) SDS-PAGE of fl-CDCP1 and c-CDCP1 (Cut 1, Cut 2, Cut 3) ectodomain show successful expression and purification. NTF is heavily glycosylated and runs as a high-molecular weight smear. (**C**) BLI of IgG 4A06 to fl- or c-CDCP1 ectodomains show that the NTF of CDCP1 is intact on both cleaved and uncleaved CDCP1. (**D**) Differential Scanning Fluorimetry (DSF) shows that fl- and c-CDCP1 have similar melting profiles and stabilities, suggesting the NTF/CTF complex does not dissociate until full unfolding of the protein. T_m_ is reported as an average and standard deviation of two replicates. (**E**) Circular Dichroism (CD) spectra of fl- and c-CDCP1. CDCP1 has a β-sheet signature with minima ∼217 nm. The slight difference in spectral shape between fl- and c-CDCP1 indicates a subtle change in secondary structure. (**F**) SAXS-derived P(r) function of fl- and c-CDCP1 ectodomains show similar overall architecture. SAXS-derived *ab initio* envelopes are shown under the graph. (**G**) Radii of gyration (R_g_) of fl- and c-CDCP1. (**H**) SEC-MALS chromatograms of fl- and c-CDCP1 show similar elution profiles and molecular weights corresponding to monomeric ectodomain.

### Overexpression of both cleaved and uncleaved CDCP1 induces downstream signaling

To study the function of the cleaved CDCP1 complex, we designed a set of lentiviral vectors to generate stable HEK293T cell lines expressing fl- or c-CDCP1 (**Fig. 4A**). For fl-CDCP1, a vector encoding the entire CDCP1 protein sequence was used. For c-CDCP1, we designed a vector in which a T2A self-cleaving sequence is placed between the CTF and the NTF. The T2A sequence is cleaved during translation to generate two polypeptides (*39*), enabling cell surface expression of the c-CDCP1 complex from a single vector. We also generated cell lines expressing fl- or c-CDCP1 variants where the four intracellular tyrosines were mutated to phenylalanine one by one (Y707F, Y734F, Y743F, Y806F) or altogether (4YF: Y707F/Y734F/Y743F/Y806F) (*40*). Flow cytometry (**Fig. 4B**) and western blot using (**Fig. 4C**) confirmed the successful generation of these stable cell lines.

**Figure 4:**
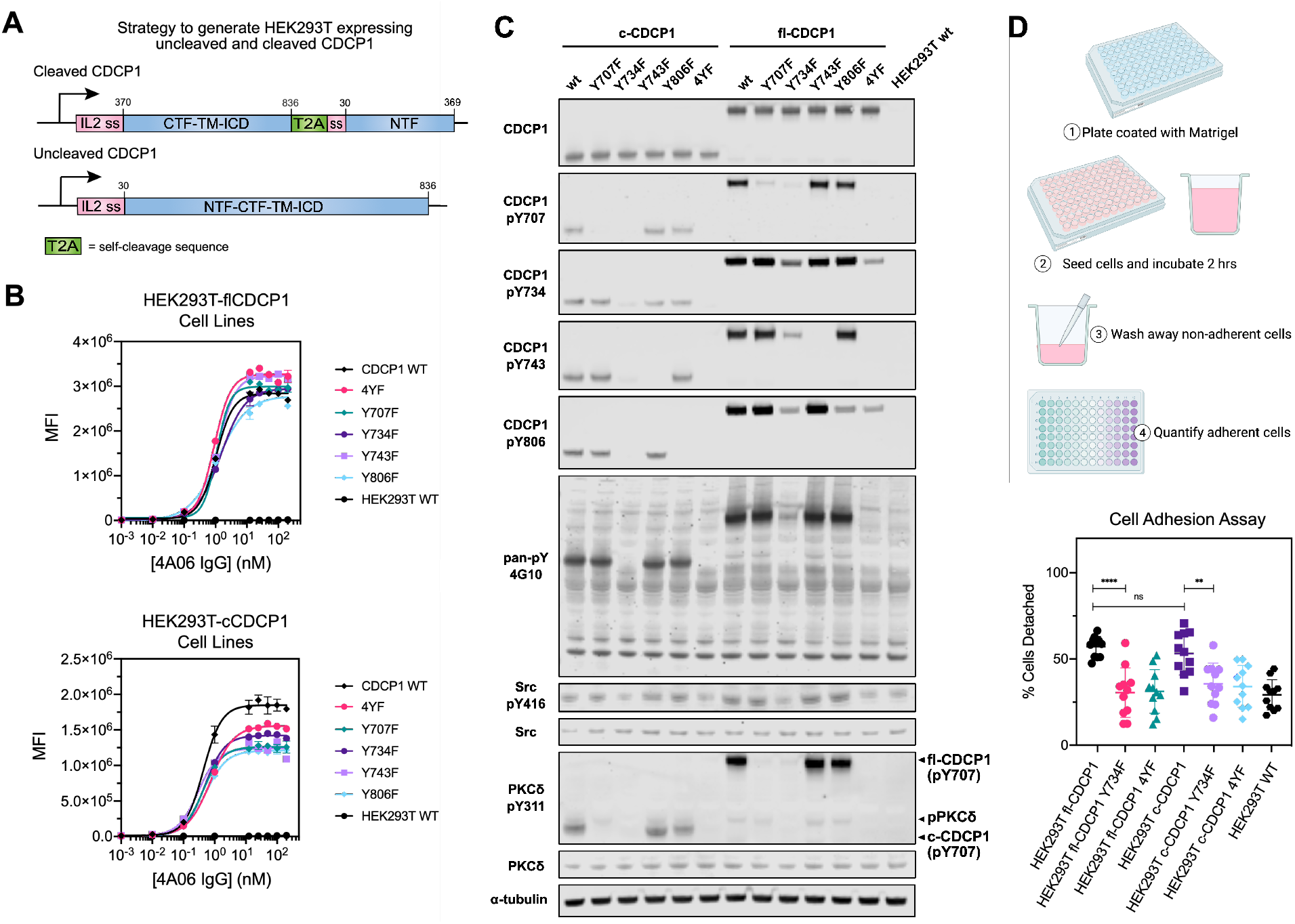
Both fl- and c-CDCP1 induce signaling and promote loss of adhesion. (**A**) Schematic of strategy to generate HEK293T cell lines expressing fl- or c-CDCP1. For c-CDCP1, a lentiviral vector was designed where a T2A self-cleavage sequence flanks the CTF (res 370-836) and NTF (res 30-369). For fl-CDCP1, a lentiviral vector encoding the full CDCP1 sequence (res 30-836) was designed. An IL2 signal sequence precedes each fragment. (**B**) Flow cytometry of IgG 4A06 to HEK293T fl-CDCP1 and HEK293T c-CDCP1 cell lines indicates the NTF of CDCP1 is present on the cell surface for all cell lines. (**D**) Western blot of CDCP1 and intracellular proteins associated with CDCP1 signaling. Both fl-CDCP1 and c-CDCP1 are phosphorylated and initiate downstream signaling mediated by Src and PKCδ. Phosphorylation of Y734 on CDCP1 is important for phosphorylation of other tyrosine residues and downstream signaling partners. Note, anti-phosphoY311-PKCδ appears to be cross-reactive to CDCP1-pY734. (**D**) Cell adhesion assay comparing HEK239T fl-CDCP1 and c-CDCP1 shows that overexpression of both fl- and c- CDCP1 decreases cell adhesion and is dependent on phosphorylation of intracellular tyrosine residues, specifically of Y734. **p = 0.0016, ****p < 0.0001, ns = not significant p >0.05. (unpaired t-test)

Overexpression of CDCP1 is associated with intracellular tyrosine phosphorylation and initiation of signaling pathways involving Src and PKCδ to promote pro-tumorigenic processes such as loss of adhesion and anoikis (*24, 40*). Western blot of CDCP1, Src, and PKCδ show that intracellular tyrosine residues of both fl- and c-CDCP1 can be phosphorylated (**Fig. 4C**) and leads to phosphorylation of Src and PKCδ. Additionally, we find that consistent with literature, Y734 is important for the phosphorylation of the other intracellular tyrosine residues of CDCP1 and the phosphorylation of Src and PKCδ (*40*). We then examined how CDCP1 expression affects cell growth and adhesion. Both overexpression of fl- and c-CDCP1 decreased cell adhesion and was dependent on intracellular tyrosine phosphorylation, specifically of Y734 (**Fig. 4D, Fig. S6**). We find that CDCP1 overexpression did not have a significant effect on cell growth (**Fig. S7)**. The expression levels of fl- and c-CDCP1 are not the same—c-CDCP1 expression is lower than fl- CDCP1, likely due to the low efficiency of the T2A self-cleaving sequence, which prevents the direct comparison of phenotypic differences between uncleaved and cleaved CDCP1. Regardless, our results show that both forms of CDCP1, if overexpressed, can be phosphorylated and initiate signaling pathways and cellular phenotypes consistent with known CDCP1 biology.

### IgG CL03 specifically recognizes the cleaved form of CDCP1

To generate an antibody that can specifically recognize cleaved CDCP1, we employed a differential phage selection strategy using an in-house Fab-phage library (**Fig. 5A**) (*41*). Prior to each round of selection, the phage pool was cleared with fl-CDCP1-Fc before positive selection with c-CDCP1-Fc. Purified antigens containing the three different cut sites were selected for individually, or as a pooled c-CDCP1 antigen mix. After 3-4 rounds of selection, there was enrichment for Fab-phage that bound c-CDCP1-Fc over fl-CDCP1-Fc (**Fig. S8A**). We identified a unique clone, CL03, which bound all three c-CDCP1 antigens selectively over fl-CDCP1 with sub-nanomolar IgG affinity (K_D_ = 150-840 pM) (**Fig. 5B, Fig. S8B, Table S2**). Additionally, we found that plasmin-treated CDCP1 can also be recognized by IgG CL03 (**Fig. S9A-C**). Since plasmin is reported to be one of the proteases that can cleave CDCP1 (*33, 42*), this suggests that IgG CL03 is capable of recognizing a physiologically relevant epitope on cleaved CDCP1.

**Figure 5:**
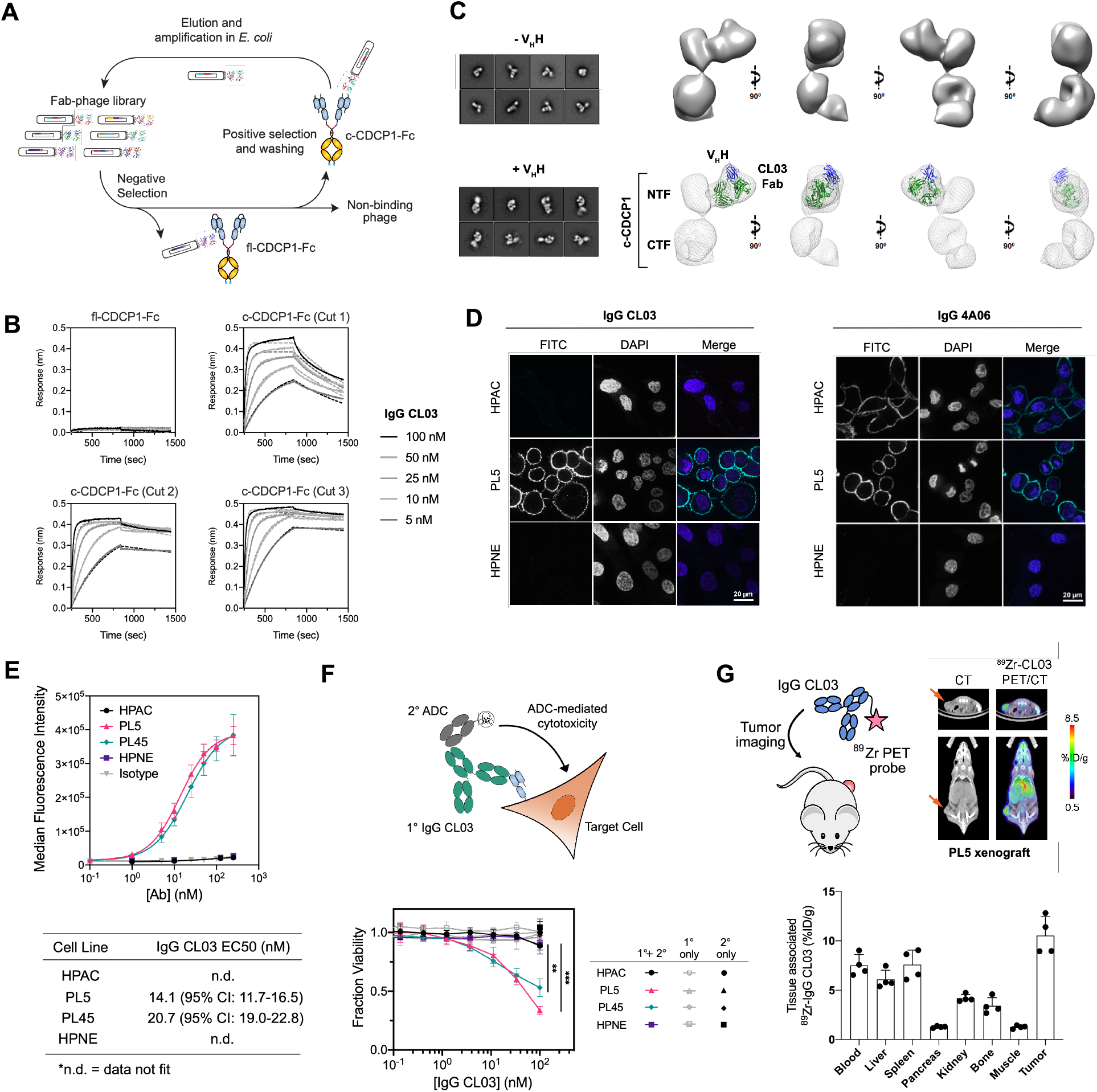
IgG CL03 specifically targets cleaved CDCP1-expressing pancreatic cancer cells. (**A**) Differential phage selection strategy to identify a cleaved CDCP1-specific antibody. Fab-phage were pre-cleared with fl-CDCP1-Fc prior to positive selection with c-CDCP1-Fc. Enriched Fab-phage were characterized by phage ELISA for selective binding to c-CDCP1-Fc. (**B**) BLI show specific binding of IgG CL03 to c-CDCP1-Fc but not to fl-CDCP1-Fc. (K_D_ = 150-840 pM, **Table S1**) (**C**) Negative-stain EM 3D reconstruction of c-CDCP1 with CL03 Fab. (*left*) 2D class averages of c-CDCP1(Cut3) + CL03 Fab in the absence and presence of anti-Fab V_H_H. (*right*) Different views of 3D EM map of CDCP1(Cut3) + CL03 Fab + V_H_H with crystal structure of Fab (green) and V_H_H (blue) modeled into the density. (**D**) Immunofluorescence of Alexa Fluor-488-labeled IgG CL03 (*left panels*) and IgG 4A06 (*right panels*) on HPAC, PL5, and HPNE cells. IgG CL03 specifically stains PL5 cells that express cleaved CDCP1, while IgG 4A06 stains both HPAC and PL5 cells. (**E**) Flow cytometry shows that IgG CL03 binds to cleaved CDCP1-expressing PL5 and PL45 cells but not HPAC or HPNE cells. (n = 3, data represent average and standard deviation). (**F**) (*top*) Schematic of antibody drug conjugate (ADC) cell killing assay. (*bottom*) Dose-dependent ADC-mediated cell killing with IgG CL03 was only observed against PL5 and PL45 cells that express cleaved CDCP1, and only in the presence of both the primary and secondary antibody. (**p = 0.004, ***p = 0.0018, unpaired T-test) (**G**) *In vivo* PET imaging of ^89^Zr-labeled IgG CL03 in PDAC xenograft mice harboring PL5 tumors (n = 4).

We were interested in understanding how CL03 is able to differentiate between fl- and c-CDCP1. One possible mechanism is that CL03 would recognize the cleavage “scar” which could include the new N- and C-termini, though this could be challenging with a heterogeneous cut site. Alternatively, CL03 could recognize an epitope that is unmasked upon proteolysis or recognize a proteolysis-induced conformational change. To investigate this, we tested the binding of CL03 to the three different c-CDCP1 antigens by BLI. We found that if the C-termini of the NTF was immobilized, either via an Fc domain or directly to the biosensor surface, there was no binding of CL03 (**Fig. S10A-B**). However, if the NTF was immobilized via the N-termini, CL03 bound NTF similarly to c-CDCP1 (**Fig. S10B-C**). This suggests that the CL03 epitope is located on the NTF, and a free C-termini is important for recognition. Interestingly, we found that CL03 can also recognize an *uncleaved* CDCP1 variant where a 16-amino acid linker is inserted between CUB1 and CUB2 at the R368/K369 site (**Fig. S10D**). This indicates that akin to proteolysis, extending the linker between CUB1 and CUB2 can also unmask the CL03 epitope.

We obtained a 3D reconstruction of c-CDCP1 (Cut 3) ectodomain bound to CL03 Fab at 25 Å resolution (**Fig. 5C, Fig. S11B-C**) and bound to 4A06 Fab at 23 Å resolution (**Fig. S11A, Fig. S11D-E**) by negative-stain electron microscopy. A nanobody that binds at the “elbow” of the light chain was used to determine the orientation and “handedness” of the Fab (*43*). No structure of CDCP1 has been reported previously. c-CDCP1 adopts an elongated structure with three distinct “lobes” of density, which likely corresponds to the three CUB domains. We reasoned from our BLI data that the Fab-bound domain is the NTF, and the other lobes belong to the CTF. CL03 appears to bind the NTF at its side, while 4A06 binds at the apex of the NTF at a distinct, non-overlapping epitope. Taken together, we propose a model in which the epitope of CL03 is located on the NTF but is inaccessible in the uncleaved form, and proteolysis releases the C-termini of NTF to unmask this epitope (**Fig. S10E**). This could be achieved by rearrangement of the secondary structure elements of cleaved CDCP1, even while adopting a compact overall conformation similar to the uncleaved form.

### IgG CL03 targets cleaved CDCP1-expressing PDAC cells

We proceeded to test whether IgG CL03 can specifically recognize and target cleaved CDCP1 on cancer cells. IgG CL03 stains cleaved CDCP1-expressing PL5 and PL45 cells with EC50 values of 14.1 and 20.7 nM, respectively, with no detectable binding to HPAC or HPNE (**Fig. 5D, 5E**). We were able to increase IgG CL03 binding to HPAC by treatment with plasmin, a serine protease known to cleave CDCP1 (**Fig. S9D-E**). We then tested whether an antibody-drug-conjugate (ADC) strategy could be used to specifically deliver cytotoxic payloads to cleaved CDCP1-expressing PDAC cells. HPAC, PL5, PL45, and HPNE cells were treated with IgG CL03 as the primary Ab along with a secondary antibody conjugated to cytotoxin monomethyl auristatin F (MMAF) (**Fig. 5F**). We observed dose-dependent cell killing of only PL5 and PL45 cells, while HPAC and HPNE were spared. Next, we tested whether CL03 as a bi-specific T-cell engager (BiTE) can selectively recruit and activate immune cells in the presence of cleaved CDCP1-expressing target cells. We genetically fused Fab CL03 to an anti-CD3 OKT3 scFv and tested whether this BiTE molecule could activate a NFAT-GFP reporter Jurkat cell line in co-culture with PDAC cells (**Fig. S12**). We observed Jurkat activation in co-culture with PL5 and PL45 cells, while co-culture with HPAC and HPNE resulted in only baseline activation. Lastly, we tested whether radiolabeled IgG CL03 can localize to cleaved CDCP1-expressing tumors in a mouse xenograft model. ^89^Zr-labeled IgG CL03 was injected into mice harboring subcutaneous PL5 tumors. Position emission tomography (PET) imaging 48 hrs post-injection showed strong tumor localization of ^89^Zr-IgG CL03 (**Fig. 5G**). Taken together, these studies demonstrate that IgG CL03 is capable of targeting cleaved CDCP1-expressing pancreatic cancer cells in a variety of modalities.

### IgG58, a cleaved-specific antibody to the mouse homolog of CDCP1, demonstrates anti-tumor activity with enhanced safety profile in a syngeneic mouse model

CL03 is cross-reactive to cynomolgous c-CDCP1 but not to the mouse homolog (**Fig. S13**). To enable syngeneic studies, we utilized the same differential phage display selection strategy against mouse CDCP1 to identify binders that specifically recognize mouse c-CDCP1 (**Fig. S14A-C**). After characterization and affinity maturation, we arrived at a lead mouse cleaved-specific CDCP1 antibody, IgG58, which binds mouse c-CDCP1 with high affinity and specificity (**Fig. 6A, Table S3**). In parallel, we identified IgG12, which, akin to the human CDCP1-specific IgG 4A06, recognizes both mouse fl- and c- CDCP1 with similar affinities (**Fig. S14D, Table S3**). We also generated a stable mouse c-CDCP1 cell line using the same T2A self-cleavage sequence strategy described previously (**Fig. 4A**) in the background of Fc1245, an aggressive mouse KPC cell line (*44*). Both IgG58 and IgG12 recognizes Fc1245 c-CDCP1 cells, binding with an EC50 of 6.9 nM and 0.46 nM, respectively (**Fig. 6B, Fig. S14E**). We further show that IgG58 and IgG12, when re-formatted to BiTE molecules, can activate Jurkat cells in the presence of Fc1245 c-CDCP1 cells (**Fig. S14F-G**). Additionally, we directly conjugated MMAF to IgG58 and IgG12 and show that these ADC molecules can specifically deliver cytotoxic payloads to Fc1245 c-CDCP1 cells while sparing Fc1245 WT cells that do not express CDCP1 (**Fig. 6C, Fig S14H**).

**Figure 6:**
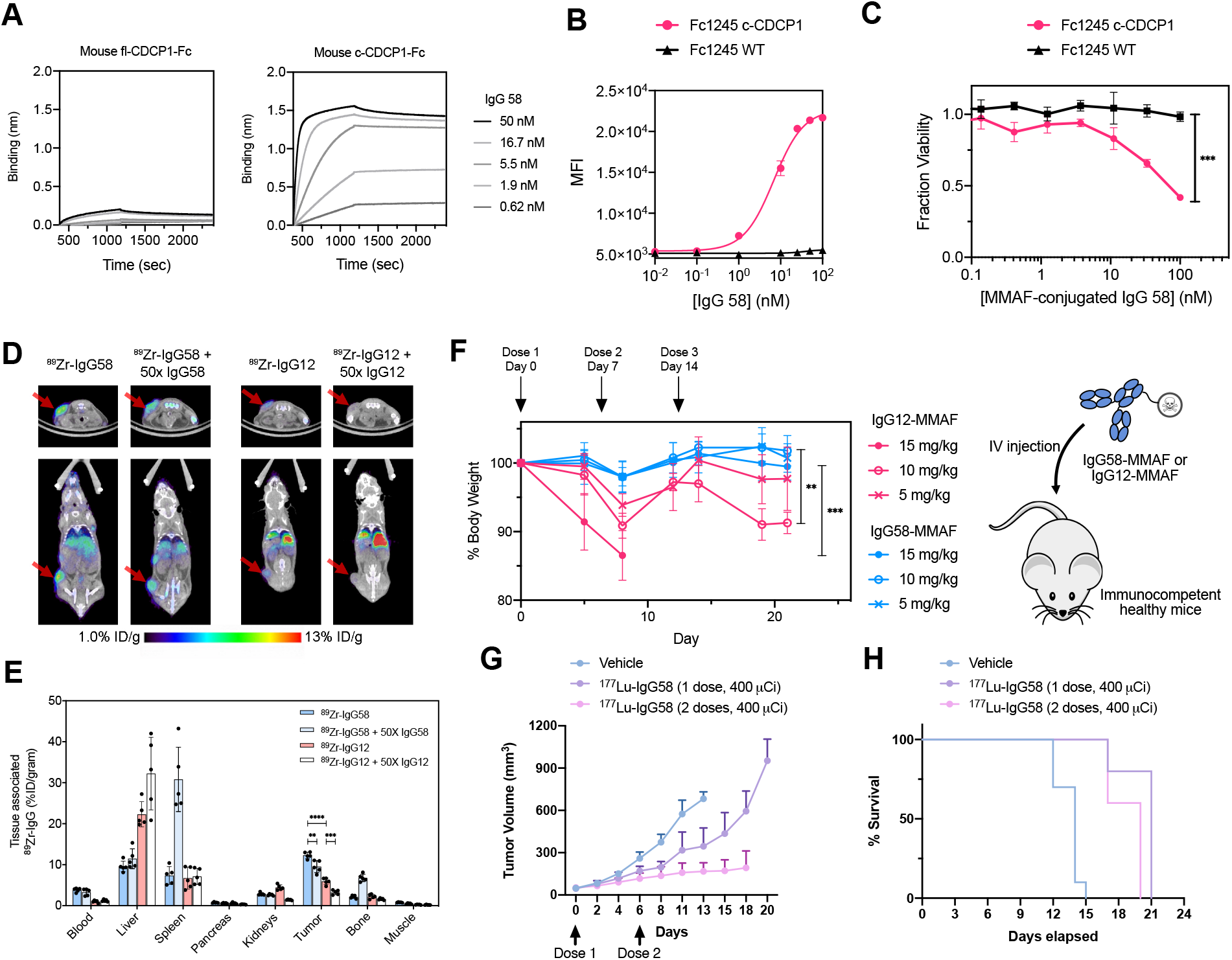
IgG58, a mouse cleaved CDCP1-specific antibody, targets cleaved CDCP1-expressing cancer cells in a mouse syngeneic pancreatic tumor model. (**A**) BLI show specific binding of IgG58 to mouse c-CDCP1-Fc, but not to fl-CDCP1-Fc. (**B**) Flow cytometry shows that IgG58 binds robustly to Fc1245 c-CDCP1, but not to Fc1245 WT cells (n = 3, error bars represent s.d.). (**C**) Dose-dependent ADC-mediated cell killing with IgG58-MMAF treatment was only observed with Fc1245 c-CDCP1 cells and not Fc1245 WT cells. (n = 2, error bars represent s.d.) (**p = 0.002, unpaired T-test) (**D**) Representative *in vivo* PET images of ^89^Zr-IgG58 and ^89^Zr-IgG12 in mice harboring subcutaneous Fc1245 c-CDCP1 tumors. Co-administration of 50X unlabeled IgG was used to examine target specificity. (**E**) Biodistribution of ^89^Zr-IgG58 and ^89^Zr-IgG12 in mice harboring subcutaneous Fc1245 c-CDCP1 tumors (n = 5 per arm). Both ^89^Zr-IgG58 and ^89^Zr-IgG12 signal decreased when 50X unlabeled IgG was administered, indicating target-specific localization (**p = 0.003, ***p = 0.0002, unpaired T-test). ^89^Zr-IgG58 shows stronger signal in the tumor, while ^89^Zr-IgG12 shows weaker tumor localization and more widespread normal tissue distribution (****p <0.0001, unpaired T-test). (**F**) ADC toxicity assay in non-tumor bearing mice. Mice (n = 5 per arm) were dosed weekly with 5, 10, 15 mg/kg of either IgG12-MMAF or IgG58-MMAF and body weight was monitored for treatment-associated toxicity. There was a significant difference between the treatment arms (F(5,32) = 3.11, p = 0.0002, ANOVA), with IgG58-MMAF treatment being better tolerated, with significant differences between IgG12-MMAF and IgG58-MMAF treatments at the 15 mg/kg dose (***p = 0.0068) and 10 mg/kg dose (**p = 0.0067) (Tukey’s multiple comparisons test). (**G-H**) Theranostic efficacy study of ^177^Lu-IgG58. Mice (n = 5 per treatment arm, n = 8 for vehicle arm) were injected with 400 µCi of ^177^Lu-IgG58 or vehicle 4 days after Fc1245-c-CDCP1 tumor implantation. For the 2-dose arm, the treatment was repeated 6 days later. Treatment with ^177^Lu-IgG58 resulted in decreased tumor growth and increased survival compared to the vehicle arm (***p = 0.0008, ****p <0.0001, unpaired two-tailed T-test).

We then tested whether IgG12 and IgG58 can localize to Fc1245 c-CDCP1 tumors *in vivo*. ^89^Zr-labeled IgG12 or IgG58 was injected into mice harboring a subcutaneous Fc1245 c-CDCP1 tumor and examined 48 hrs later by PET imaging (**Fig. 6D, 6E**). High tumor localization of ^89^Zr-IgG58 was observed. The signal decreased when 50X unlabeled IgG58 was co-administered, indicating tumor-specific localization driven by target engagement. We observed minimal ^89^Zr-IgG58 signal systemically. In comparison, we observe weaker tumor-specific localization of ^89^Zr-IgG12 and more widespread off-tumor signal, suggesting the higher presence of uncleaved CDCP1 in non-tumor tissues.

We proceeded to examine the toxicity profile of MMAF-labeled IgG12 or IgG58. Non-tumor-bearing mice were dosed weekly with 5, 10, or 15 mg/kg of MMAF-labeled IgG12 and IgG58, and their body weight was monitored for 21 days. None of the mice injected with IgG58-MMAF at the 3 different doses exhibited significant changes in body weight (**Fig. 6F**). In contrast, mice treated with IgG12-MMAF experienced significant body weight loss following the administration of each dose, indicative of treatment-induced toxicity. All of the mice receiving the 15 mg/kg dose of IgG12-MMAF had to be euthanized due to body weight loss by Day 8, and 2 of the 5 mice receiving the 10 mg/kg dose of IgG12-MMAF were euthanized on Day 19 for the same reasons. These results suggest that a cleaved CDCP1-targeting molecule would have a superior safety profile compared to a pan-CDCP1-targeting approach.

Lastly, we investigated the anti-tumor activity of IgG58 as a radioligand therapeutic. Treating mice harboring subcutaneous Fc1245 c-CDCP1 tumors with one or two 400 µCi doses of ^177^Lu-IgG58 significantly reduced tumor volume compared with mice receiving vehicle control, with the two-dose regimen approaching tumor stasis (**Fig. 6G, Fig. S15**). Median survival for the treatment arms were 21 and 20 days for the 1- and 2-dose regimen, respectively, compared to 14 days for the vehicle group. A significant survival advantage was imparted by ^177^Lu-IgG58 theranostic therapy (**Fig. 6H**), supporting our conclusion that antibodies specific to proteolytic neo-epitopes could expand the targetable disease space for cancer treatment.

## DISCUSSION

CDCP1 was first identified as a highly upregulated gene in colorectal and lung cancer (*18*) and has emerged as a potential therapeutic target and a driver of tumorigenesis and metastasis across a wide range of indications (*16, 23–25, 30, 45–49*). The functional role of CDCP1 has been linked to well-established oncogenic signaling networks including Ras (*21, 29*), EGFR (*50*), PDGFR (*47*), HER2 (*51*), and HIF (*52*). Interest in the therapeutic potential of CDCP1 is demonstrated by numerous studies that have developed small molecules (*53*) and antibodies (*18, 21, 26, 41, 54, 60, 61*) against to CDCP1 and its pathways. However, despite these efforts, an anti-CDCP1 therapeutic has yet to enter the clinic. Although CDCP1 is highly expressed on cancer cells at nearly ∼2 million copies per cell (*21, 22*), it is also present on normal epithelial tissue (*55*). There is evidence that CDCP1 is not cleaved during normal physiological processes, but its cleavage is induced during tumorigenesis (*56*). This and other reports (*34–36*) suggest that cleaved CDCP1 would be rare on the surface of normal cells. Our work demonstrates that selectively targeting the cleaved isoform of CDCP1 could be a safer, more therapeutically attractive approach. Continued work to characterize the prevalence and role of cleaved CDCP1, particularly in clinically relevant samples will help determine which patient populations and disease types would be best suited to a cleaved-CDCP1 targeting strategy.

Interestingly there is remarkably little conformational change between cleaved and uncleaved CDCP1. This presents a new model for CDCP1 proteolysis and has been recently corroborated by Kryza et al. (*57*) where they also observe that the N-terminal fragment of CDCP1 does not dissociate upon proteolysis. We determined the exact sites of proteolysis and utilized multiple biophysical and biochemical methods on both recombinant and cell surface CDCP1 to further bolster the evidence that cleaved CDCP1 is a complex and has similar conformation to the uncleaved form. Despite this, we were able to raise antibodies that are highly selective to the cleaved form by rigorous positive/negative selections. We reason that proteolysis between CUB1 and CUB2 exposes an epitope on the NTF that can be recognized by CL03. It is possible, however, that there are conditions in which the NTF dissociates from the membrane-bound fragment, as shed CDCP1 can be detected in the serum of cancer patients (*58*).

We observe that both uncleaved and cleaved CDCP1, when overexpressed, can be phosphorylated and induce downstream signaling and associated cellular phenotypes. This is not surprising given that SEC, EM, and SAXS all show little difference in conformation between cleaved and uncleaved CDCP1. However, it has been shown, in some cases, that proteolysis leads to increased CDCP1 phosphorylation and signaling (*27, 31, 33*). Because the expression levels of fl-CDCP1 and c-CDCP1 were not the same in our study, we are not able to assess whether proteolysis differentially affects fl-CDCP1 vs c-CDCP1 signaling (*24*). CDCP1 can also be processed by multiple serine proteases (*28, 33*), interact with other membrane proteins to form signaling complexes (*27, 49*), and is regulated by other cancer-associated programs (*50, 59–61*), all of which adds complexity to CDCP1’s role in normal and tumor biology. Future studies that continue to dive into the biology of CDCP1 could illuminate whether proteolysis drives function or whether it is a surrogate marker for elevated proteolysis on the surface of cancer cells.

Extracellular neo-epitopes generated by alternate splice forms (*62*), MHC-peptide complexes (*63*), post-translational modifications (PTMs) (*64*), and glycosylation (*65*) are emerging as an attractive class of therapeutic targets for cancer and other disease. However, characterizing these neo-epitopes and developing therapeutic molecules that can selectively recognize them are challenging. Complications include but are not limited to the low-abundance of MHC-peptide complexes, rarity of alternative splice variants that present targetable epitopes, the heterogeneity of glycosylation, and finding and validating proteoforms that are truly disease-specific. Proteolysis is irreversible, highly prevalent in the tumor microenvironment, and can alter the conformation and structure of proteins and protein complexes in both subtle and significant ways. We hypothesized that proteolysis-generated neo-epitopes on the cancer cell surface could provide an alternative, orthogonal approach to expand the therapeutic index of targeted therapeutics. The cleaved-specific CDCP1 antibodies described here demonstrate that specifically targeting cleaved-CDCP1 is effective and has a more favorable safety profile compared to targeting pan-CDCP1. We believe this is a unique demonstration of an antibody specific to a proteolytically processed form of a cancer-associated cell surface protein. Given the important and widespread role of proteases in disease biology, proteolysis-induced neo-epitopes are likely to be present on other proteins (*8*) and could be targeted with antibodies using this same rigorous positive/negative selection strategy.

## MATERIALS AND METHODS

### Cloning, protein expression, and purification

Plasmids encoding CDCP1 as Fc or non-Fc fusions, the heavy chain and light chains of IgGs, and the heavy chain and light chain fused to scFv of BiTE were generated by Gibson cloning into pFUSE (InvivoGen). Fabs were PCR-ed from the Fab phagemid and subcloned into pBL347. Plasmids for stable cell line construction were generated by Gibson cloning into pCDH-EF1-CymR-T2A-Neo (System Bioscience). Sequences of all plasmids were confirmed by Sanger sequencing.

We used a previously described vector for expression of Fabs.^66^ Briefly, C43 (DE3) Pro+ E. coli transformed with the Fab expression plasmid were grown in TB autoinduction media at 37°C for 6 hrs, then switched to 30°C for 16–18 hrs. Cells were harvested by centrifugation (6000xg for 20 min) and lysed with B-PER Bacterial Protein Extraction Reagent (Thermo Fischer). Lysate was incubated at 60°C for 20 min and centrifuged at 14,000xg for 30 min. Clarified supernatant was passed through a 0.45 µm syringe filter. Fabs were purified by Protein A affinity chromatography on an AKTA Pure system. Fab purity and integrity were assessed by SDS-PAGE and intact mass spectrometry.

Constructs encoding CDCP1 were generated by transfection of BirA-Expi293 cells (Life Technologies) with one or two plasmids encoding cleaved or uncleaved CDCP1 antigen as Fc or non-Fc fusions. IgGs and BiTE were generated by transfection of BirA-Expi293 cells with two plasmids encoding the heavy chain and light chain at 1:1 ratio. The ExpiFectamine 293 transfection kit (Life Technologies) was used for transfections as per manufacturer’s instructions. Cells were incubated for 5 days at 37°C in 5% CO2 at 125 rpm before the supernatants were harvested by centrifugation. Proteins were purified by Protein A affinity chromatography (CDCP1 Fc-fusions, IgGs, and BiTE) or Ni-NTA affinity chromatography (non-Fc fusion CDCP1) and assessed for quality and integrity by SDS-PAGE.

### Lentiviral cell line construction

All the HEK293T stable cell lines were generated by lentiviral transduction. To produce virus, HEK293T Lenti-X cells were transfected with a mixture of second-generation lentiviral packaging plasmids at ∼80% confluence. FuGene HD (Promega) was used for transfection of the plasmids using 3 µg DNA (1.35 µg pCMV delta8.91, 0.15 µg pMD2-G, 1.5 µg pCDH vectors encoding gene of interest) and 7.5 µL of FuGene HD per well of a six-well plate. Media was changed to complete DMEM after 6 hrs of incubation with transfection mixture. The supernatant was harvested and cleared by passing through a 0.45 µm filter 72 hrs post transfection. Cleared supernatant was added to target HEK293T wild-type cells (∼1 million cells per mL) with 8 µg/mL polybrene and cells were centrifuged at 1000xg at 33°C for 2 hrs. Cells were then incubated with viral supernatant mixture overnight before the media was changed to fresh complete DMEM. Cells were expanded for a minimum of 48 hrs before they were grown in drug selection media. Drug selection for stable cell lines was started by the addition of 2 µg/mL puromycin. Following at least 72 hrs of incubation in puromycin containing media, cells were analyzed by flow cytometry for expression of CDCP1.

Mouse pancreatic cancer cell line Fc1245 overexpressing c-CDCP1 were generated using the same protocol except using a Hygromycin B selection marker.

### Mammalian Cell Culture

HPAC, PL5, PL45, and HPNE cells were a gift from the laboratory of E. Scott Seeley (Stanford) and were maintained in IMDM + 10% FBS + 1X Pen/Strep. The HEK293T cell lines were cultured in DMEM + 10% FBS + 1X Pen/Strep. Jurkat NFAT-GFP reporter cells lines were cultured in RPMI + 10% FBS + 2 mg/mL G418 + 1X Pen/Strep. Fc1245 was cultured in DMEM + 10% FBS + 1X Pen/Strep. Cell line identities were authenticated by morphological inspection. Symptoms for mycoplasma contamination were not observed and thus no test for mycoplasma contamination was performed. To the best of our knowledge, all cell lines that were received as gifts were previously authenticated and tested for mycoplasma.

### Immunoprecipitation (IP)

Cells confluent in a 15-cm dish were washed by PBS buffer three times before being lysed on plate with 1 mL ice-cold NP-40 lysis buffer supplemented with protease inhibitor cocktail (Roche) and PhosSTOP (Roche). The cell lysate was incubated at 4°C with gentle shaking for 30 min and centrifuged at 14,000xg for 30 min at 4°C. The supernatant was then transferred into a new tube. Protein concentration was determined by BCA assay (Bio-Rad). For IP-WB, 5 µL CDCP1 antibody (D1W9N, Cell Signaling) was added into 200 µL cell lysate precleared by 20 µL Protein A magnetic beads (EMD Millipore) and incubated overnight at 4°C. On the second day, 20 µL Protein A magnetic beads was added to the lysate and incubated for 20 min at rt. The Protein A magnetic beads were washed 5x with 0.5 mL lysis buffer. 20 µL 4X SDS loading buffer was added and the beads were heated to elute protein at 95°C for 5 min. The sample was then analyzed by WB. For IP-MS, 20 µL CDCP1 antibody (D1W9N, Cell Signaling) was added into 1 mL cell lysate and incubated overnight at 4°C. On the second day, 100 µL of Protein A magnetic beads was added to the lysate and incubated for 20 min. Protein A magnetic beads were washed 5x with 1 mL lysis buffer and PBS buffer. CDCP1 was then eluted with 200 µL 0.1 M acetic acid and neutralized by 20 µL pH 11 Tris Buffer. Samples were then used for proteomics analysis.

### Mass Spectrometry (MS)

Samples were reduced with 5 mM TCEP at 55°C for 30 min and alkylated with 10 mM iodoacetamide at room temperature for 30 min. Proteins were subjected to digestion using 20 µg sequencing grade Glu-C (Promega) in 1 M urea at 37°C overnight. The eluted fraction was collected and then desalted using Sola column (Thermo Fisher) following standard protocol. Desalted peptides were dried and dissolved in mass spectrometry buffer (0.1% formic acid + 2% acetonitrile) prior to LC-MS/MS analysis.

1 µg of peptide was injected into a pre-packed 0.075 mm x 150 mm Acclaim Pepmap C18 LC column (2 µm pore size, Thermo Fisher) attached to a Q Exactive Plus (Thermo Fisher) mass spectrometer. Peptides were separated using a linear gradient of 3–35% solvent B (Solvent A: 0.1% formic acid, Solvent B: 80% acetonitrile, 0.1% formic acid) over 170 min at 300 µL/min. Data were collected in data-dependent acquisition mode using a top 20 method with a dynamic exclusion of 35 sec and a charge exclusion restricted to charges of 2, 3, or 4. Full (MS1) scan spectrums were collected as profile data with a resolution of 140,000 (at 200 m/z), AGC target of 3E6, maximum injection time of 120 msec, and scan range of 400–1800 m/z. Fragment ion (MS2) scans were collected as centroid data with a resolution of 17,500 (at 200 m/z), AGC target of 5E4, maximum injection time of 60 msec with a normalized collision energy at 27, and an isolation window of 1.5 m/z with an isolation offset of 0.5 m/z. Peptide search and MS1 peak area quantification were performed using ProteinProspector (v.5.13.2) against user-defined CDCP1 sequence (Uniprot Accession ID: Q9H5V8).

### Western Blot

The confluent cells were washed with PBS and lysed with ice-cold NP-40 Lysis Buffer supplemented with protease inhibitor cocktail (Roche) and PhosSTOP (Roche). Lysate was incubated with gentle shaking at 4°C for 30 min and centrifuged at 14,000xg for 30 min at 4°C. The supernatant was then transferred into a new tube and protein concentration was determined by BCA assay (Bio-Rad). Immunoblotting was performed using CDCP1(D1W9N) (Cell Signaling, 13794S), Phospho-CDCP1(Tyr707) (Cell Signaling, 13111S), Phospho-CDCP1(Tyr806) (Cell Signaling, 13024S), Phospho-CDCP1(Try734) (Cell Signaling, 9050S), Phospho-CDCP1(Try743)(D2G2J) (Cell Signaling, 14965S), Src(36D10) (Cell Signaling, 2109S), Phospho Src Family (Tyr416) (Cell Signaling, 2101S), PKCδ (Cell Signaling, 2058S), Phospho-PKCdelta(Tyr311) (Cell Signaling, 2055S), alpha-Tubulin(DM1A) (Cell Signaling, 3873S), IRDye 680RD Goat anti-Mouse (LiCOR, 925–68070), and IRDye 800CW Goat anti-Rabbit (LiCOR, 926-32211) antibodies.

### Flow cytometry

Cells were lifted with Versene (0.04% EDTA, PBS pH 7.4 Mg/Ca free), washed once with PBS pH 7.4, and subsequently blocked with flow cytometry buffer (PBS, pH 7.4, 3% BSA). Primary Abs were added to cells for 30 minutes at 4°C. Bound Abs were detected with addition of AlexaFluor-488 or AlexaFluor-647 conjugated Goat anti-human IgG, F(ab’)2 fragment specific (Jackson ImmunoResearch: 1:1000). Cells were washed 3x with PBS + 3% BSA and fluorescence was quantified using a CytoFLEX (Beckman Coulter) flow cytometer. All flow cytometry data analysis was performed using FlowJo software and Prism software (GraphPad).

### Bio-layer interferometry (BLI) experiments

BLI experiments were performed using an Octet RED384 instrument (ForteBio). Biotinylated proteins were immobilized on a streptavidin (SA) biosensor and His-tagged proteins were immobilized on a Ni-NTA bisoensor. Different concentrations of analyte in PBS pH 7.4 + 0.05% Tween-20 + 0.2% BSA (PBSTB) were used as analyte. Affinities (KDs) were calculated from a global fit (1:1) of the data using the Octet RED384 software.

### Size Exclusion Chromatography (SEC)

SEC analysis was performed using an Agilent HPLC 1260 Infinity II LC System using an AdvanceBio SEC column (300 Å, 2.7 μm, Agilent). Each analyte was injected at 1-10 μM and run with a constant mobile phase of 0.15 M sodium phosphate for 15 min. Absorbance at 280 nm was measured.

### Circular Dichroism (CD) Spectroscopy

CD spectra were measured using an Aviv 410 CD spectrophotometer. The CD signal from 200 nm to 300 nm was collected in a 0.1-cm path length cuvette at 25°C. Samples contained 0.4 mg/mL of protein in PBS. All CD spectra were blanked with PBS in the absence of protein.

### SEC-SAXS

Small angle X-ray scattering in-line with size exclusion chromatography (SEC-SAXS) data were collected at the SIBLYS beamline 12.3.1 of the Advanced Light Source at the Lawrence Berkeley National Laboratory^67^ at 1.127 Å wavelength and sample-to-detector distance of 2,105 mm. The resulting scattering vectors, defined as *q* = 4π sinθ/λ, where 2θ is the scattering angle, ranged from 0.01 to 0.4 Å^−1^. Data were collected using a Dectris PILATUS3 2M detector at 20°C and processed as previous described.^68,69^

The SEC-SAXS flow cell was directly coupled with an Agilent 1260 Infinity HPLC system using a Shodex KW-803 column. The column was equilibrated with running buffer (1X PBS - 10 mM phosphate, 137 mM NaCl, 2.7 mM KCl; pH 7.4) at a flow rate of 0.45 mL/min. 50 µL of fl-CDCP1 or c-CDCP1 (Cut 3) proteins were injected at ∼ 5 mg/ml and 3 sec X-ray exposures were collected continuously over the 30 min SEC elution in the running buffer for each sample. The SAXS frames recorded prior to the protein elution peak were used to subtract the signal for SAXS frames across the elution peak. Radius of gyration (R_g_) were determined based on the Guinier approximation,^70^ I(q) = I(0) exp (-q^2^R_g_^2^/3), with the limits qR_g_ < 1.3. Further, scattering intensity at q = 0 Å^−1^ (I(0)) and R_g_ values were compared for each collected SAXS curve across the entire elution peak. The elution peak was mapped by comparing the integral of ratios to background and Rg relative to the recorded frame using the program SCÅTTER. Interference-free SAXS curves with least R_g_ variation were averaged and merged in SCÅTTER to produce the highest signal-to-noise SAXS curves. These merged SAXS curves were used to generate the Guinier plots, volumes-of-correlation (V_c_), pair distribution functions, P(r), and normalized Kratky plots. Pair distribution function P(r) was computed using program GNOM^71^ and use to determine the maximal dimension of the macromolecule. P(r) functions were normalized based on the molecular weight (MW) of the macromolecules, as determined based on their calculated V_c_.^72^

### SEC-MALS data collection and analysis

The SEC eluent was split (4 to 1) between the SAXS line and a multiple wavelength detector (UV-vis) at 280 nm, multi-angle light scattering (MALS), and refractometer. MALS experiments were collected using an 18-angle DAWN HELEOS II light scattering detector connected in tandem to an Optilab refractive index concentration detector (Wyatt Technology). Bovine serum albumin was used for system normalization and calibration. The MALS data complimented the MW determined by SAXS analyses. The MALS data were also used to align the SAXS and UV-vis peaks along the X-axis (elution volume in mL) to compensate for fluctuations in timing and band broadening. MALS and differential refractive index data were analyzed using Wyatt Astra seven software to monitor the homogeneity of the sample molecular weights across the elution peak.

### Differential Scanning Fluorimetry (DSF)

Protein in PBS were mixed with Sypro Orange dye (20 X stock) to make a final protein concentration of 2 µM and dye concentration 4X. 10 µL of each mixture was transferred to a Biorad 384-well PCR white plate and the plate was covered by qPCR Sealing Tape. The assay was performed over temperature range of 25°C to 95°C with a temperature ramping rate of 0.5°C/30 seconds on a Roche LC480 Light Cycler.

### Negative-stain Electron Microscopy

Complexes of c-CDCP1 with CL03 Fab and 4A06 Fab (in a molar ratio 1:1) in the absence or presence of a V_H_H antibody were obtained by size exclusion chromatography on Superdex 200 increase 10/300 GL column in the buffer containing 10 mM HEPES, 100 mM NaCl, pH 7.5. 5 µL of protein sample at 0.006 mg/mL (for c-CDCP1-4A06 +/- V_H_H) or 0.008 mg/mL (for c-CDCP1- CL03 +/- V_H_H) was applied on a glow-discharged formvar/carbon-coated TEM grid and negatively stained using a sequential four-droplet method. Each grid with the sample was washed in two 50 µL drops of distilled water for one second and two 50 µL drops of 1% uranyl formate solution for one and thirty seconds, respectively. After removing excess of uranyl formate solution, grids were dried at room temperature. Grids were screened on a FEI Tecnai G2 F30 300 kV Super Twin TEM Electron Microscope (FEI Company, Hillsboro, OR) at Advanced Electron Microscopy Facility at the University of Chicago (Chicago, IL). Micrographs were acquired at magnification 49 kX with a pixel size of 0.23 nm on the level of specimen using a 4K x 4K CCD camera. Particles were selected automatically using RELION.^73^ Extracted particles were 2D class averaged, sorted into initial classes, 3D classified and refined in RELION. Final maps were analyzed in UCSF Chimera.^74^

### Cell Proliferation Assay

Cell proliferation assays were performed using an MTT modified assay to measure cell viability. In brief, 5,000 HEK293T cells expressing different CDCP1 constructs were plated in each well of a 96-well plate on day 0. Cells were incubated at 37°C under 5% CO2. On day 1, 2, 3 and 4, 10 µL of 5 mg/mL of Thiazolyl Blue Tetrazolium Bromide (Sigma Aldrich) was added to each well and incubated at 37°C under 5% CO2 for 2 hrs. Following, 100 µL of 10% SDS + 0.01 M HCl was added to lyse the cells to dissolve the MTT product. After 4 hrs, absorbance at 595 nm was quantified using an Infinite M200 PRO-plate reader (Tecan). Data points were plotted using Prism software (GraphPad).

### Cell Adhesion Assay

The day prior to the assay, a 96-well tissue culture plate was coated with MaxGel™ ECM (Millipore Sigma) 1:10 diluted in serum-free DMEM. At the same time, the cell culture medium for HEK293T cells with expressing different CDCP1 constructs was changed to serum-free medium. The next day, media was removed and the culture plates were blocked with 100 μL serum-free DMEM with 0.1 % BSA for 2 hrs and washed with PBS. 100,000 cells in 100 μL of serum-free (0.1 % BSA) medium were added to each well and incubated at 37°C in a 5% CO2 for 2 hrs. The non-adherent cells were removed by washing with media three times, and the remaining cells were quantified by MTT assay. Data points were plotted using Prism software (GraphPad).

### Phage selection

All phage selections were done according to previously established protocols.^41^ Briefly, selections were performed using biotinylated c-CDCP1-Fc captured on SA-coated magnetic beads (Promega). Prior to each selection, the phage pool was incubated with 1 µM of biotinylated fl-CDCP1-Fc captured on streptavidin beads in order to deplete the library of any binders to fl-CDCP1. Four rounds of selection were performed with decreasing amounts of c-CDCP1-Fc (100 nM, 50 nM, 10 nM and 10 nM). We employed a “catch and release” strategy, where bound Fab-phage were eluted from the magnetic beads by the addition of 2 µg/mL of TEV protease. Individual phage clones from the third and fourth round of selection were analyzed for binding by phage ELISA.

### Phage ELISA

Phage ELISAs were performed according to standard protocols. Briefly, 384-well Maxisorp plates were coated with NeutrAvidin (10 μg/mL) overnight at 4°C and subsequently blocked with PBS + 2% BSA for 1 hr at 20°C. 20 nM of biotinylated c-CDCP1-Fc or fl-CDCP1 ECD-Fc was captured on the NeutrAvidin-coated wells for 30 min followed by the addition phage supernatants diluted 1:5 in PBSTB for 30 min. Bound phage were detected using a horseradish peroxidase (HRP)-conjugated anti-M13 phage antibody (GE Lifesciences 27-9421-01).

### Immunofluorescence

HPAC, PL5, and HPNE cells were plated on glass-bottom imaging plates (MatTek) and incubated for 24 hrs at 37°C under 5% CO2. Cells were treated with IgG (1 µg/mL) for 30 min and washed with media to remove unbound IgG. Bound IgG was detected by the addition of a Alexa Fluor® 488-conjugated AffiniPure F(ab’)2 Fragment Goat Anti-Human IgG, F(ab’)2 Fragment Specific (Jackson ImmunoResearch, 143225) in Invitrogen Molecular Probes Live Cell Imaging Solution (Thermo Fisher) containing Hoescht blue (2 µg/mL). Cells were imaged on a Nikon Ti Microscope Yokogawa CSU-22 with Spinning Disk Confocal.

### Antibody Drug Conjugate (ADC) Labeling

The ADC labeling protocol was adapted from previous reports.^75,76^ To 2 mL IgG58 or IgG12 in PBS buffer with 50 mM borate pH 8.0 was added DTT to a final concentration of 10 mM. After incubation at 37 °C for 30 min, the buffer was exchanged by elution through G-25 column with PBS containing 1 mM diethylenetriamine pentaacetic acid (DTPA, sigma). PBS containing 1 mM DTPA (PBS/D) was added to the reduced antibody to make the antibody concentration 2.5 mg/mL in the final reaction mixture, and the solution was chilled. The drug-linker (9 e.q. of antibody, MC-Val-Cit-PAB-MMAF, MedChemExpress) solution to be used in the conjugation was prepared by diluting drug-linker from a frozen dimethyl sulfoxide (DMSO) stock solution at a known concentration (approximately 10 mM) in sufficient DMSO to make the conjugation reaction mixture 5-10% organic/95-90% aqueous, and the solution was chilled on ice. The drug-linker solution was added rapidly with mixing to the cold-reduced antibody solution, and the mixture was left on ice for 1 hour. A 20-fold excess of cysteine over maleimide was then added from a freshly prepared 100-mM solution in PBS/D to quench the conjugation reaction. While the temperature was maintained at 4°C, the reaction mixture was concentrated by centrifugal ultrafiltration and buffer-exchanged by elution through Sephadex G25 equilibrated in PBS. The conjugate was then filtered through a 0.2-μm filter under sterile conditions and stored at –80°C for analysis and testing.

### In vitro Antibody Drug Conjugate (ADC) Assays

*In vitro* antibody drug conjugate cell killing assays were performed either using a secondary antibody conjugated to MMAF, or by direct conjugation of MMAF to the primary antibody. Secondary ADC assays used a Fab Anti-Human IgG Fc-MMAF Antibody with Cleavable Linker (Moradec) as a secondary antibody following manufacture’s protocol. For ADC assays using direct conjugation of MMAF to the primary antibody, the antibody was labeled with DBCO-PEG4-ValCit-MMAF (Levena Biosciences) site-specifically at residue T74M of the light chain using oxazirdine chemistry using a previously described protocol.^77^ A day prior to the assay, 5000 cells were seeded on a 96-well poly-lysine-coated white plate (Corning). The next day, media was removed and 100 µL of primary IgG and secondary ADC at a 1:4 ratio or 100 µL of MMAF-labeled primary IgG in media was added. Cells were incubated for 72 hrs at 37°C under 5% CO_2_. After the incubation period, 100 µL of CellTiter-Glo Reagent (Promega) was added to each well followed by incubation at room temperature for 10 min with gentle shaking. Luminescence was measured using an Infinite M200 PRO plate reader (Tecan).

### Bi-specific T-cell engager (BiTE) Assay

A day prior to the assay, 25,000 target cells were seeded on a 96-well plate (Corning). The next day, media was removed and 50,000 Jurkat NFAT-GFP reporter cells and BiTE (10-fold dilution) were added to a final media volume of 200 µL. After incubation for 20 hrs at 37°C, Jurkat cells were recovered by gentle pipetting, washed in PBS + 3% BSA, and GFP expression was quantified by flow cytometry using a CytoFLEX (Beckman Coulter) flow cytometer.

### Mouse Positron Emission Tomography (PET) Imaging Study

#### Nucleotide conjugation for PET imaging

250 µL of IgG (1 mg) at a concentration of 4.0 mg/mL was mixed with 150 µL of 0.1 M sodium bicarbonate buffer (pH 9.0). The pH was adjusted to 9.0 and the final reaction mixture was adjusted to a total volume of 0.5 mL by adding a sufficient amount of 0.1 M sodium bicarbonate buffer. Df-Bz-NCS (p-isothiocyanatobenzyl-desferrioxamine) was dissolved in DMSO at a concentration of 10 mM. Df-Bz-NCS solution was added to the antibody solution to give a three-molar excess of the chelator over the molar amount of IgG. The Df-Bz-NCS was added in steps of 2 µL and mix rigorously during the addition. The concentration of DMSO was kept below 2% of the total reaction mixture in order to avoid any precipitation. After 60 min at 37°C, the reaction mixture was purified via a G-25 column pre equilibrated by 9 mL of PBS (pH 7.4). The IgG-DFO solution was eluted in PBS and aliquoted for radiolabeling.

In a reaction vial, 89Zr-oxalic acid solution (5mCi; 10 µL) was neutralized with 200uL of 1 M HEPES (pH 7.4) After 1 minute, 0.5 mg of IgG-DFO (pH 7.4) was then added into the reaction vial. After incubation (60 min) at 37°C, the radiolabeling efficiency was determined by ITLC using chromatography strips and 20 mM citric acid (pH 4.9–5.1). The radiolabeling efficiency was consistently >98.5%. An additional purification and buffer exchange step were performed with a G-25 column pre-equilibrated with PBS. The 89Zr-IgG was eluted in PBS and further diluted with 0.9% Sodium Chloride Injection, USP, before being administered into mice for PET imaging.

#### Tumor implantation

Mouse PET imaging studies were conducted in compliance with Institutional Animal Care and Use Committee at UCSF. IgG CL03 PET imaging was conducted on nu/nu mice bearing subcutaneous human pancreatic PL5 and HPAC xenograft tumors. IgG 58 or IgG 12 PET imaging was conducted on syngeneic mouse model of immunocompetent Black6(C57Bl/6J) mice bearing subcutaneous mouse pancreatic Fc1245 c-CDCP1 tumors. 4 to 6 weeks-old intact male athymic nu/nu mice were purchased from Charles River Laboratory and healthy male Black6(C57BL/6J) mice of 6-8 weeks were purchased from the Jackson Laboratory. All the mice were used for experiments after a brief period of acclimation. Mice were inoculated subcutaneously with PL5 (∼5.0 × 106), HPAC (∼5.0 × 106) or Fc1245 c-CDCP1 (∼1.0 × 106) cells in the flank. The cells were injected in a 1:1 mixture (v/v) of media (IMDM for PL5 and HPAC, DMEM for Fc1245) and Matrigel (Corning). Imaging studies were conducted after the tumors reached an appropriate size.

#### Small-animal PET/CT

All data, including the raw imaging files, are available upon request. Tumor-bearing mice received between approximately 200 μCi of 89Zr-IgG CL03, 12 or 58 in 100 μL saline solution volume intravenously using a custom mouse tail vein catheter with a 28-gauge needle and a 100–150 mm long polyethylene microtubing. After a dedicated period of uptake time (48 hours), mice were anesthetized with isoflurane and imaged on a small-animal PET/CT scanner (Inveon, Siemens Healthcare). Animals were scanned for 20 minutes for PET, and the CT acquisition was performed for 10 minutes. The coregistration between PET and CT images was obtained using the rigid transformation matrix generated prior to the imaging data acquisition because the geometry between PET and CT remained constant for each of PET/CT scans using the combined PET/CT scanner. The photon attenuation was corrected for PET reconstruction using the coregistered CT-based attenuation map to ensure the quantitative accuracy of the reconstructed PET data. Decay corrected images were analyzed using AMIDE software.

#### Biodistribution studies

At the dedicated time points after radiotracer injection, animals were euthanized by cervical dislocation. Blood was harvested via cardiac puncture. Tissues were removed, weighed, and counted on a Hidex automatic gamma counter for accumulation activity. The mass of the injected radiotracer was measured and used to determine the total number of counts per minute (CPM) by comparison with a standard of known activity. The data were background and decay corrected and expressed as the percentage of the injected dose/weight of the biospecimen in grams (%ID/g).

#### ADC Toxicity Study

The ADC toxicity was performed with healthy male Black6(C57BL/6J) mice of 8-10 weeks old purchased from the Jackson Laboratory. Mice (n = 5 in each group) were dosed intravenously weekly for 3 weeks with ADCs (IgG58-MMAF and IgG12-MMAF at 15, 10, 5 mg/kg in 200 µL PBS buffer). Body weight was monitored biweekly for 4 weeks total. Mice were sacrificed if body weight dropped below 80% of original. All experiments were performed in accordance with relevant guidelines and regulations and in full accordance with UCSF Institutional Animal Care and Use Committee. Statistical analysis was performed using a one-way ANOVA in GraphPad Prism.

### Antibody–Nucleotide Conjugate (ANC) Efficacy Study

#### Nucleotide conjugation for efficacy study

200 µL of IgG 58 (2 mg) at a concentration of 10.0 mg/mL was mixed with 200 µL of 0.1 M sodium bicarbonate buffer (pH 9.0). The pH was adjusted to 9.0 and the final reaction mixture was adjusted to a total volume of 0.5 mL by adding a sufficient amount of 0.1 M sodium bicarbonate buffer. p-SCN-Bn-DOTA (S-2-(4-Isothiocyanatobenzyl)-1,4,7,10-tetraazacyclododecane tetraacetic acid) was dissolved in DMSO at a concentration of 50 mM. p-SCN-Bn-DOTA solution was added to the antibody solution to give a fifty-molar excess of the chelator over the molar amount of IgG. The p-SCN-Bn-DOTA was added in steps of 2 µL and mix rigorously during the addition. The concentration of DMSO was kept at no more than 2% of the total reaction mixture in order to avoid any precipitation. After 90 min at 37°C, the reaction mixture was purified via a G-25 column pre equilibrated by 9 mL of 0.2 M Ammonium Acetate (pH 7.0). The IgG58-DOTA solution was eluted in 0.2 M Ammonium acetate and aliquoted for radiolabeling.

In a reaction vial, 177Lu-chloride solution (6 mCi; 3 µL) was added directly to 900 uL (2 mg) of IgG58-DOTA (pH 7.0). After incubation (60 min) at 37°C, the radiolabeling efficiency was determined by ITLC using chromatography strips and 20 mM citric acid (pH 4.9–5.1). The radiolabeling efficiency was consistently >98.5%. An additional purification and buffer exchange step was performed with a PD-10 column pre-equilibrated with PBS. The 177Lu-IgG 58 was eluted in PBS and further diluted with 0.9% Sodium Chloride Injection, USP, before being administered into mice for ANC treatment.

#### Tumor implantation

Mouse ANC efficacy studies were conducted in compliance with Institutional Animal Care and Use Committee at UCSF. Healthy male Black6(C57BL/6J) mice of 4-6 weeks were purchased from the Jackson Laboratory and used for experiments after a brief period of acclimation. Mice were inoculated subcutaneously with mouse pancreatic cancer cells (Fc1245 c-CDCP1 ∼ 1.0 × 106) in the flank. The cells were injected in a 1:1 mixture (v/v) of media (DMEM) and Matrigel (Corning). Treatments were started 3 days post tumor implantation.

#### ANC treatment

Mice bearing unilateral subcutaneous Fc1245 c-CDCP1 tumors received 177Lu-IgG 58, or vehicle (saline) at the indicated dose via tail vein. Mice were weighed and tumor sizes were measured at the time of injection, and three times weekly until the completion of the study. Tumor volume measurements were calculated with calipers. For treatment studies, the primary endpoints were death due to tumor volume >2,000 mm3, tumor ulceration, or ≥20% loss in mouse body weight.

## Supplementary Materials

Fig. S1: BLI of IgG 4A06 binding to the N-terminal fragment (NTF) of CDCP1.

Fig. S2: Thrombin treatment of CDCP1 ectodomain with engineered thrombin cut site generates cleaved CDCP1 complex.

Fig. S3: CDCP1 peptides identified by IP-MS of PDAC cell lines.

Fig. S4: Generation of fl-CDCP1 and c-CDCP1 ectodomain with endogenous cut sites.

Fig. S5: SEC-SAXS of uncleaved and cleaved CDCP1 ectodomain show similar conformations.

Fig. S6: Cell Adhesion Assay of HEK293T cells expressing CDCP1 tyrosine variants show that phosphorylation of Y734 is critical for detachment.

Fig. S7: MTT Cell proliferation assay of HEK293T cell lines expressing fl-CDCP1 and c-CDCP1 variants show over-expression, phosphorylation, or cleavage have no effect on cell proliferation.

Fig. S8: Identification of cleaved CDCP1-specific Fab by phage selection.

Fig. S9: IgG CL03 recognizes plasmin-cleaved CDCP1.

Fig. S10: CL03 binds to an epitope on the NTF that is exposed on cleaved CDCP1, but not uncleaved CDCP1.

Fig. S11: Negative stain EM 3D reconstruction of cleaved CDCP1 bound to 4A06 Fab.

Fig. S12: CL03 as a Bi-specific T-cell engager (BiTE) can activate Jurkat cells in the presence of cleaved CDCP1-expressing PDAC cells.

Fig. S13: Species cross-reactivity of IgG CL03.

Fig. S14: Targeting mouse CDCP1 with IgG12 and IgG58.

Fig. S15: Tumor volume of individual mice in ^177^Lu-IgG58 theranostic study.

Table S1: SAXS and MALS experimental parameters

Table S2: In vitro binding affinities of Fab CL03 and IgG CL03 to uncleaved and cleaved CDCP1

Table S3: Binding affinity of mouse CDCP1 antibodies to cleaved and uncleaved forms of mCDCP1

## Acknowledgements

We thank members of the Wells Lab for helpful discussion and input, in addition to members of the Recombinant Antibody Network for feedback on this project. We would also like to thank Dr. Dan Zhu, Dr. Hari Hariharan, Dr. Henry Chan, and Dr. Ho Cho of Bristol Myers Squibb for their insights into this work. We also thank Dr. Shalini Chopra, Nima Hooshdaran, Fernando Salangsang, and Paul Phojanakong for assistance in the *in vivo* mouse studies. We also thank Dr. Susan Marqusee (UC Berkeley) for access to a CD spectrophotometer, and Dr. Michal Hammel and Daniel Rosenberg of BL12.3.1 for their assistance in data collection and initial analysis.

## Author contributions

SAL and JZ designed and conducted all experiments unless otherwise noted. AJM conducted the characterization of Prescission-cleavable CDCP1. YW and MJE conducted or supervised the PET imaging experiments. EVF and AAK conducted or supervised the EM data collection and analysis. VS, DW, BH conducted or supervised the mouse experiments. SGR prepared and analyzed the SAXS experiments. JL conducted the DSF experiments. KKL and JAW supervised the research. SAL, JZ, JAW prepared and wrote the manuscript with input from all authors.

## Data and materials availability

All data are available in the main text or the supplementary materials.

