## Supplementary material for "Targeting a proteolytic neo-epitope of CUB-domain containing protein 1 in RAS-driven cancer": SI

**This PDF file includes:**

Materials and Methods

Figs. S1 to S15

Tables S1 to S3

**Fig. S1: BLI of IgG 4A06 binding to the N-terminal fragment (NTF) of CDCP1.** NTF, composed of residues 30-367 of CDCP1, fused to a C-terminally biotinylated Fc domain was immobilized on Streptavidin-coated biosensors and probed with 25 nM of IgG 4A06. IgG 4A06 binds robustly to NTF-Fc, indicating that 4A06 recognizes an epitope within the NTF of the CDCP1 ectodomain.

**Fig. S2: Thrombin treatment of CDCP1 ectodomain with engineered thrombin cut site generates cleaved CDCP1 complex.** (**a**) SDS-PAGE gel of Thrombin protease-cleavable CDCP1-Fc (CDCP1(Tx)-Fc). The reported R368/K369 cleavage site was replaced with a Thrombin protease recognition sequence (GS)_5_-LVPRGS-(GS)_5_. Treatment with Thrombin protease cleaves CDCP1(Tx)-Fc between the CUB1 and CUB2 domains and generate the corresponding NTF and CTF-Fc fragments. (**b**) BLI of IgG 4A06 binding to C-terminally immobilized CDCP1(Tx)-Fc treated or untreated with Thrombin shows that IgG 4A06, that recognizes the NTF, can still bind cleaved CDCP1. (**c**) SEC of CDCP1(Tx)-Fc treated or untreated with Thrombin protease shows that cleaved and uncleaved CDCP1 have similar elution profiles suggesting the NTF and CTF-Fc remain associated.

**Fig. S3: CDCP1 peptides identified by IP-MS of PDAC cell lines.** Peptides identified by IP-MS of 3 different PDAC cell lines (a) PL5, (b) PL45, (c) HPAC. Peptides corresponding to cleaved fragments of CDCP1 were identified on PL5 and PL45 cells, but not HPAC.

**Fig. S4: Generation of fl-CDCP1 and c-CDCP1 ectodomain with endogenous cut sites.** (**a**) Schematic of a two-plasmid co-transfection strategy to generate the cleaved CDCP1 ectodomain as an Fc fusion. The NTF and CTF-Fc were encoded on separate plasmids, with an IL2 signal sequence preceding each sequence. (**b**) SDS-PAGE gel of fl-CDCP1-Fc and c-CDCP1-Fc (Cut 1, Cut 2, Cut 3) (**c**) BLI shows robust binding of IgG 4A06 to fl-CDCP1-Fc and c-CDCP1-Fc. (**c**) BLI of IgG 4A06 to fl-CDCP1-Fc and c-CDCP1-Fc (Cut 1, Cut 2, Cut 3) show robust binding, indicating NTF is present on all 4 constructs. (**d**) SEC of fl-CDCP1-Fc and c-CDCP1-Fc (Cut 1, Cut 2, Cut 3) shows that all 4 antigens have similar elution profiles. (**e**) SEC of fl-CDCP1 and c-CDCP1 (Cut 1, Cut 2, Cut 3) without Fc domains shows that all 4 antigens have similar elution profiles.

**Fig. S5: SEC-SAXS of uncleaved and cleaved CDCP1 ectodomain show similar conformations.** (**a**) SAXS profiles, (**b**) Normalized Kratky plot, and (**c**) P(r) function of fl-CDCP1 and c-CDCP1 (Cut 3) ectodomain indicates that there are no large-scale conformational changes between fl-CDCP1 and c-CDCP1.

**Table S1: SAXS and MALS experimental parameters**

|  | **R_g_  (Å)** | **Dmax (Å)** | **MW MALS/SAXS (kDa)** |
| --- | --- | --- | --- |
| **fl-CDCP1** | 45.4 +/- 0.6 | 145 | 97.0 (+/- 0.11%) / ~112 |
| **c-CDCP1 (Cut3)** | 44.1 +/- 0.5 | 143 | 99.6 (+/- 0.11%) / ~108 |

**Fig. S6: Cell Adhesion Assay of HEK293T cells expressing CDCP1 tyrosine variants show that phosphorylation of Y734 is critical for detachment.** Cell adhesion assay of for HEK293T cells expressing (**a**) fl-CDCP1 and (**b**) c-CDCP1 variants. The 4 intracellular tyrosine residues (Y707, Y734, Y743, Y806) were individually or altogether (4YF) mutated to phenylalanine to abrogate signaling. Data indicate that Y734 is important for CDCP1-associated intracellular signaling implicated in decreased cell adhesion. Individual data points are shown, and bars indicate average. **p = 0.003, ****p <0.0001, ns = not significant, p > 0.05. Unpaired t-test using Prism software was used for statistical analysis.

**Fig. S7: MTT Cell proliferation assay of HEK293T cell lines expressing fl-CDCP1 and c-CDCP1 variants show over-expression, phosphorylation, or cleavage have no effect on cell proliferation.** Cell proliferation assay measured by MTT for HEK293T cells expressing (**a**) HEK293T WT, fl-CDCP1, and c-CDCP1, (**b**) fl-CDCP1 variants, and (**c**) c-CDCP1 variants. fl-CDCP1 or c-CDCP1 overexpression does not have a significant effect on cell growth compared to HEK293T WT cells. All variants have similar rates of growth over a 4-day period. Data were collected in triplicate and average and standard deviation are shown.

**Fig. S8: Identification of cleaved CDCP1-specific Fab by phage selection.** (**a**) Eluted phage from round 4 of phage selection indicate that there is enrichment for Fab-phage that recognize cleaved CDCP1 over uncleaved CDCP1. (**b**) BLI of Fab CL03 Fab binding to immobilized fl-CDCP1-Fc, c-CDCP1-Fc (Cut 1, Cut 2, Cut 3) shows that Fab CL03 selectively recognizes cleaved CDCP1 and not uncleaved CDCP1.

**Table S2: In vitro binding affinities of Fab CL03 and IgG CL03 to uncleaved and cleaved CDCP1**

**Fig. S9: IgG CL03 recognizes plasmin-cleaved CDCP1.** (**a**) SDS-PAGE gel of fl-CDCP1 ectodomain treated with 0.5 μg plasmin indicates that plasmin can cleave CDCP1 at the expected molecular weights to generate c-CDCP1. (**b**) BLI of fl-CDCP1 treated with plasmin shows that IgG CL03 can specifically recognize plasmin-cleaved CDCP1. Binding of IgG 4A06 indicates that the NTF remains associated to the CTF. (**c**) SEC of plasmin-treated CDCP1 shows similar elution profiles to untreated fl-CDCP1. (**d**) Flow cytometry of HPAC cells treated with plasmin shows a dose-dependent increase in IgG CL03 binding, indicating that plasmin treatment can generate cleaved CDCP1 on the surface of HPAC cells. (**e**) Flow cytometry of PL5 cells treated with plasmin shows no additional increase in IgG CL03 binding.

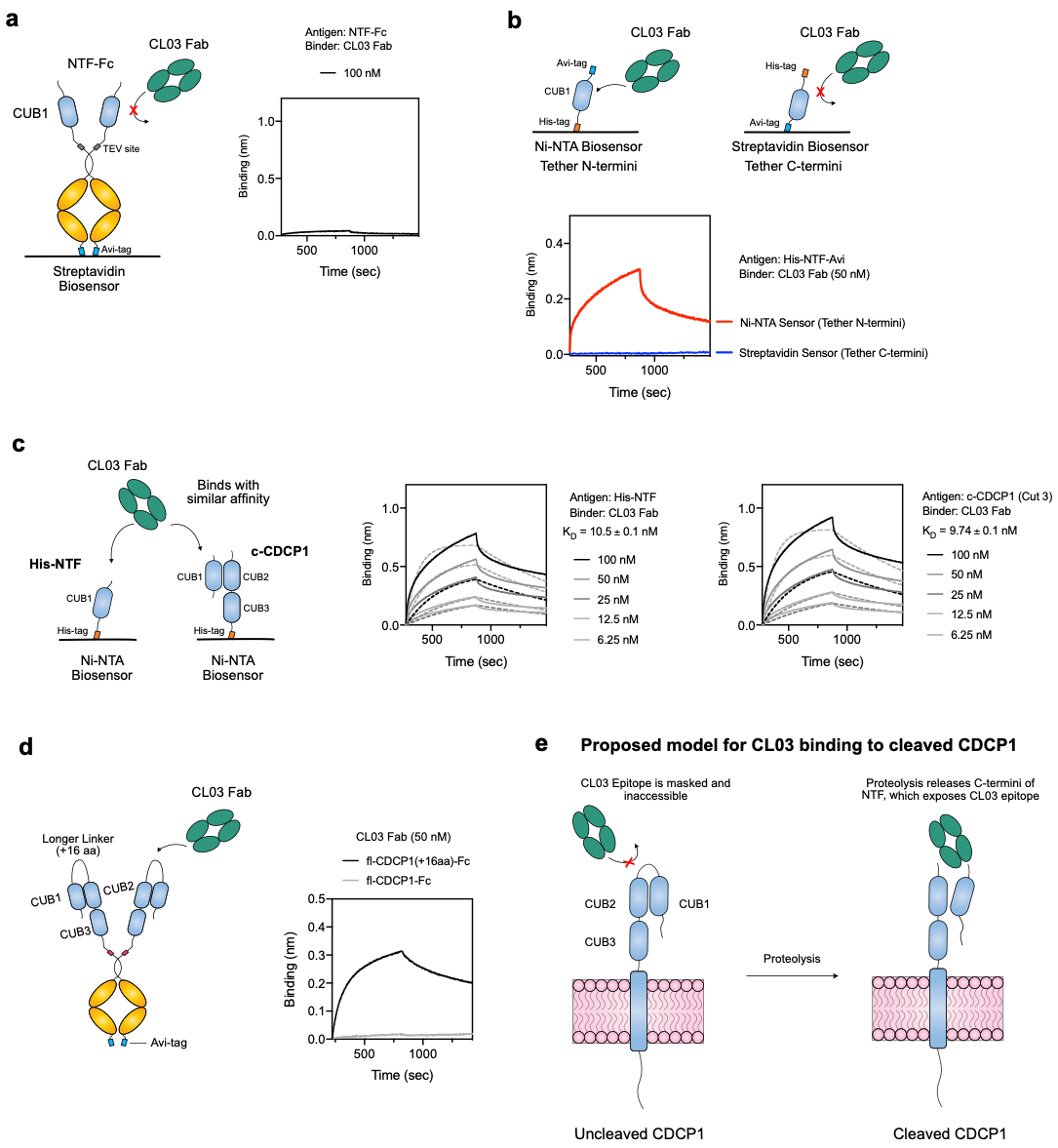

**Fig. S10: CL03 binds to an epitope on the NTF that is exposed on cleaved CDCP1, but not uncleaved CDCP1.** (**a**) BLI of CL03 Fab does not show binding to NTF-Fc. NTF-Fc was immobilized on a Streptavidin(SA) biosensor via a C-terminal biotinylated Avi-tag on the Fc domain. (**b**) BLI of CL03 Fab shows binding to N-terminally immobilized NTF, but not C-terminally immobilized NTF, indicating that an untethered C-termini of the NTF is necessary for CL03 binding. (**c**) Multipoint BLI show that CL03 Fab binds N-terminally tethered NTF at similar affinity to c-CDCP1, indicating the CL03 epitope is located within the NTF of CDCP1. (**d**) BLI shows CL03 Fab binding to uncleaved CDCP1 where the 2-residue R368/K369 cleavage site was replaced with a 16-amino acid linker. (**e**) Proposed model of CL03 binding to cleaved CDCP1. CL03 Fab recognizes an epitope on the NTF of CDCP1 that is masked and inaccessible when CDCP1 is uncleaved. Proteolysis releases the C-termini of the NTF and unmasks this epitope, allowing CL03 to bind and selectively recognize cleaved CDCP1 over uncleaved CDCP1.

**Fig. S11: Negative stain EM 3D reconstruction of cleaved CDCP1 bound to 4A06 Fab.** (**a**) (*left*) 2D class averages of c-CDCP1(Cut3) + 4A06 Fab in the absence and presence of V_H_H single domain antibody. (*right*) Different views of 3D negative stain EM map of c-CDCP1(Cut3) + 4A06 Fab + V_H_H. Crystal structure of Fab (red) with V_H_H (blue) were fitted into the 3D negative stain EM map. (**b**) A demonstrative micrograph of negatively stained c-CDCP1(Cut3) + CL03 Fab + V_H_H particles. (**c**) A Fourier shell correlation plot used to determine the model resolution of 25 Å of c-CDCP1(Cut3) + CL03 Fab + V_H_H, as given by 0.143 criterion. (**d**) A demonstrative micrograph of negatively stained c-CDCP1(Cut3) + 4A06 Fab + V_H_H particles. (**e**) A Fourier shell correlation plot used to determine the model resolution of 23 Å of c-CDCP1(Cut3) + 4A06 Fab + V_H_H, as given by 0.143 criterion.

**Fig. S12: CL03 as a Bi-specific T-cell engager (BiTE) can activate Jurkat cells in the presence of cleaved CDCP1-expressing PDAC cells.** (**a**) Schematic of bi-specific T-cell engager (BiTE)-mediated T-cell activation assay. (**b**) Dose-dependent activation of NFAT-GFP reporter Jurkat cells above background were only observed in the presence of cleaved CDCP1-expressing PL5 and PL45 cells. (n = 2, error bars represent s.d.) (**c**) BiTE CL03 (1 nM) activates NFAT-GFP reporter T-cells in the presence of PL5 and PL45 cells but not HPAC or HPNE cells. (***p<0.001, unpaired T-test).

**Fig. S13: Species cross-reactivity of IgG CL03.** BLI of IgG CL03 to (**a**) cleaved cynomolgus CDCP1 homolog and (**b**) cleaved mouse CDCP1 homolog shows that IgG CL03 is cross-reactive and selective for cynomolgus cleaved CDCP1 but does not recognize mouse uncleaved or cleaved CDCP1.

**Fig. S14: Targeting mouse CDCP1 with IgG12 and IgG58.** (**a**) Schematic of the two cut sites of cleaved mouse CDCP1. (**b**) SDS-PAGE gel of mouse CDCP1 antigens: fl-CDCP1-Fc, c-CDCP1-Fc (Cut 1), c-CDCP1-Fc (Cut 2) show proteins at the expected molecular weights. (**c**) SEC of mouse CDCP1 antigens: fl-CDCP1-Fc, c-CDCP1-Fc (Cut 1), c-CDCP1-Fc (Cut 2) show similar elution profiles, indicating the two fragments of cleaved CDCP1 is intact as a complex. (**d**) BLI of mouse CDCP1 antigens: fl-CDCP1-Fc and c-CDCP1-Fc show that IgG12, which recognizes the NTF of mouse CDCP1, can bind both cleaved and uncleaved mouse CDCP1, indicating that the NTF remains associated to the CTF. c-CDCP1-Fc was a 1:1 molar ratio of Cut 1 and Cut 2 antigens. (**e**) Flow cytometry shows that IgG58 binds robustly to Fc1245 c-CDCP1, and weakly to Fc1245 WT cells that express low levels of uncleaved CDCP1 (n = 3, error bars represent s.d.). (**f**) IgG12 reformatted to a BiTE shows dose-dependent activation of NFAT-GFP reporter Jurkat cells above background only in the presence of Fc1245 c-CDCP1 cells, but not in the presence of Fc1245 WT cells. (n = 2, error bars represent s.d.) (**g**) IgG58 reformatted to a BiTE shows dose-dependent activation of NFAT-GFP reporter Jurkat cells above background only in the presence of Fc1245 c-CDCP1 cells, but not in the presence of Fc1245 WT cells. (n = 2, error bars represent s.d.) (***p = 0.005, unpaired T-test) (**h**) Dose-dependent ADC-mediated cell killing with IgG12 and a secondary antibody conjugated to MMAF was only observed in Fc1245 c-CDCP1 cells and not Fc1245 WT cells. (n = 2, error bars represent s.d.) (**p = 0.0037, unpaired T-test)

**Table S3: Binding affinity of mouse CDCP1 antibodies to cleaved and uncleaved forms of mCDCP1**

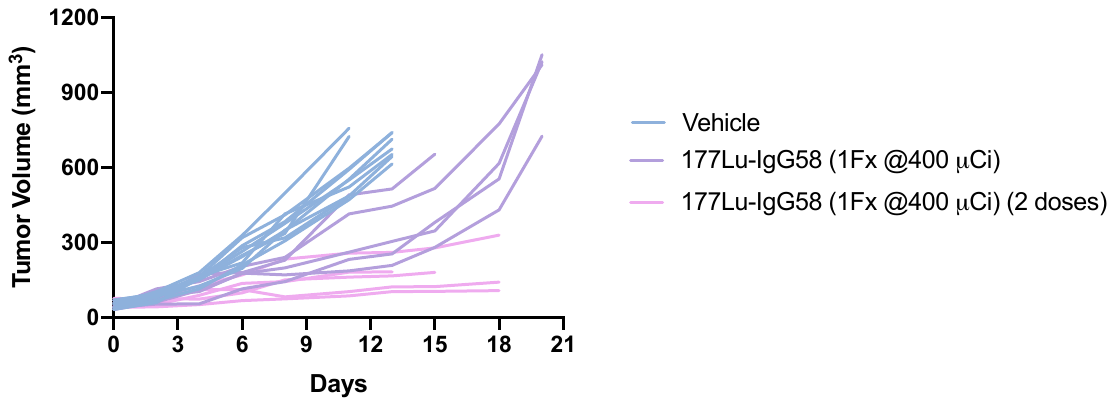

**Fig. S15: Tumor volume of individual mice in ^177^Lu-IgG58 theranostic study.** Tumor volume of mice treated with vehicle or one or two 400 µCi doses of ^177^Lu-IgG58 (n = 8 for vehicle, n = 5 per treatment arm) were monitored daily.
